# Neurons burdened by DNA double strand breaks incite microglia activation through antiviral-like signaling in neurodegeneration

**DOI:** 10.1101/2021.12.23.474002

**Authors:** Gwyneth Welch, Carles Boix, Eloi Schmauch, Jose Davila-Velderrain, Matheus B. Victor, Vishnu Dileep, Lorenzo Bozzelli, Qiao Su, Jemmie Cheng, Audrey Lee, Noelle Leary, Andreas Pfenning, Manolis Kellis, Li-Huei Tsai

**Affiliations:** Picower Institute for Learning and Memory, Massachusetts Institute of Technology, Cambridge, MA, USA; Department of Brain and Cognitive Sciences, Massachusetts Institute of Technology, Cambridge, MA, USA; Computer Science and Artificial Intelligence Laboratory, Massachusetts Institute of Technology, Cambridge, MA, USA; Broad Institute of Harvard and MIT, Cambridge, MA, USA; A. I. Virtanen Institute for Molecular Sciences, University of Eastern Finland, Kuopio, Finland; Carnegie Mellon University Departments of Computational Biology and Biology and Neuroscience Institute, Pittsburgh, PA, USA

## Abstract

DNA double strand breaks (DSBs) are linked to aging, neurodegeneration, and senescence^1,2^. However, the role played by neurons burdened with DSBs in disease-associated neuroinflammation is not well understood. Here, we isolate neurons harboring DSBs from the CK-p25 mouse model of neurodegeneration through fluorescence-activated nuclei sorting (FANS), and characterize their transcriptomes using single-nucleus, bulk, and spatial sequencing techniques. We find that neurons harboring DSBs enter a late-stage DNA damage response marked by the activation of senescent and antiviral-like immune pathways. We identify the NFkB transcription factor as a master regulator of immune gene expression in DSB-bearing neurons, and find that the expression of cytokines like *Cxcl10* and *Ccl2* develop in DSB-bearing neurons before glial cell types. Alzheimer’s Disease pathology is significantly associated with immune activation in excitatory neurons, and direct purification of DSB-bearing neurons from Alzheimer’s Disease brain tissue further validates immune gene upregulation. Spatial transcriptomics reveal that regions of brain tissue dense with DSB-bearing neurons also harbor signatures of inflammatory microglia, which is ameliorated by NFkB knock down in neurons. Inhibition of NFkB or depletion of Ccl2 and Cxcl10 in DSB-bearing neurons also reduces microglial activation in organotypic brain slice culture. In conclusion, we find that in the context of age-associated neurodegenerative disease, DSBs activate immune pathways in neurons, which in turn adopt a senescence associated secretory phenotype to elicit microglia activation. These findings highlight a novel role for neurons in the mechanism of age-associated neuroinflammation.

**Summary:** It is unclear how age-associated DNA double strand break (DSB) accumulation in neurons influences the progression of cellular senescence and neurodegenerative disease. Here, we leverage mouse models of neurodegeneration, single-nucleus, bulk, and spatial transcriptomics from Alzheimer’s disease patients, mouse models, and primary neuron cultures to dissect the immune signaling pathways initiated by DSB-bearing neurons that trigger neuroinflammation.

## Introduction

Loss of genomic integrity is linked to aging and neurodegeneration^1,2^. DNA damage repair pathways are transcriptionally prominent in the aging brain, and many age-associated neurodegenerative diseases exhibit both accumulation of DNA lesions and reduced DNA repair efficiency^3–7^. The most toxic of these lesions, the DNA double strand break (DSB), can drive many phenotypes of aging including senescence, mutation, and cell death. Postmitotic neurons are particularly susceptible to these threats due to their long lifespan, high metabolic activity, and limited DSB repair capacity. While DSB accumulation in neurons is a well-documented feature of aging and neurodegeneration, the transcriptional profile adopted by such population of neurons remains largely unknown.

The accumulation of DSBs is an early feature of Alzheimer’s Disease (AD), suggesting that they may act as an initiating lesion of toxicity^8^. Multiple mouse models of neurodegeneration phenocopy increased DSBs at early pathological stages, including the P301S tauopathy model, the inducible CK-p25 model, and the 3xTg and hAPP-J20 amyloid pathology models^9–13^. In addition, induction of DSBs are sufficient to induce organismal aging and neurodegenerative phenotypes in mice^14^. Recent studies characterizing single and double strand breaks in postmitotic neurons reveal that break location may underlie the toxicity of DNA damage in aging and neurodegenerative disease^15,16^. However, regardless of location, the downstream biological effects of DSB accumulation in neurons are unclear.

Here, we aimed to characterize the biological consequences of DSB accumulation in neurons. We also investigated how this impacts mechanisms of neuroinflammation in age-associated neurodegenerative disease. We utilized fluorescence-activated nuclei sorting (FANS) followed by bulk and single-nucleus RNA-sequencing to transcriptionally characterize neurons burdened with DSBs in the CK-p25 mouse model of neurodegeneration. We found that DSB-bearing neurons activate innate immune signaling pathways reminiscent of those expressed by senescent cells and virally infected neurons^17–24^, and that this is accompanied by degradation of neuronal identity. The gene expression patterns of DSB-burdened neurons were enriched in excitatory neurons from AD postmortem human brain. Purified DSB-bearing neurons from the human brain also exhibited an enrichment of immune gene signatures. Spatial transcriptomics of the CK-p25 forebrain revealed signatures of microglial inflammation were closely associated with DSB-bearing neurons. Correspondingly, suppression of the NFkB transcription factor in neurons impacted measures of neuroinflammation in microglia at early stages of disease. Together, these data establish a novel signaling relationship between neurons burdened with DSBs and microglia in age-associated neurodegenerative disease.

## Results

### Identification of DSB-bearing neurons at early stages of disease in a mouse model of neurodegeneration

We utilized the CK-p25 mouse model of inducible neurodegeneration to understand how DSB-bearing neurons contribute to disease development. In these mice, the *CamkII* promoter drives the expression of the neurotoxic protein fragment p25 through a doxycycline-off system^25^. P25 is the calpain-cleaved product of p35, an activator of cyclin dependent kinase 5 (Cdk5). Previously, we determined that the first pathologies observed in these mice are increased DSBs in neurons^9^ and activation of microglia^26^. These pathologies occur 1-2 weeks after the onset of p25 expression when mice are taken off doxycycline (dox). Intracellular amyloid-beta accumulation occurs as early as 2-3 weeks after induction^27^. Neuronal loss, learning deficits, and tau hyperphosphorylation are also observed in the following 4-12 weeks^25,27,28^. These pathological events are inducible and occur in a highly concerted and predictable manner^29^.

The phosphorylation of the histone variant H2A.X (γH2AX) by ATM kinase occurs rapidly after DSB detection. This post-translational histone modification is essential for efficient DSB recognition and repair, and is a robust DSB biomarker^30^. Using flow cytometry, we identified a distinct population of γH2AX-positive nuclei at the two-week time point in the CK-p25 cortex, but not in the CK control cortex (Figure 1a). CK mice express the *CamkII*-driven tetracycline transactivator (tTA), but not p25. A timeline analysis was also performed to determine when γH2AX-positive nuclei begin to accumulate in the CK-p25 cortex. γH2AX-positive nuclei were detectable as early as one week after induction (1.038 ± 0.2627 % population), peaked at two weeks (4.614 ± 0.9416 % population), and gradually decreased thereafter (Figure 1b). The significant reduction in γH2AX-positive nuclei at six weeks corresponds with previous observations of neuronal loss in this model, suggesting that γH2AX-positive cells degenerate by six weeks^25,28^. We were able to observe similar γH2AX-positive population dynamics using immunofluorescent microscopy (Supplementary Figure 1a,b). We did not detect γH2AX-positive nuclei after only four days off dox, suggesting that one week is the earliest time point at which this population appears. To validate DNA damage response pathways were active in these nuclei, we confirmed co-immunoreactivity for phosphorylated ATM kinase (pATM) (Supplementary Figure 1c,d).

**Figure 1.**
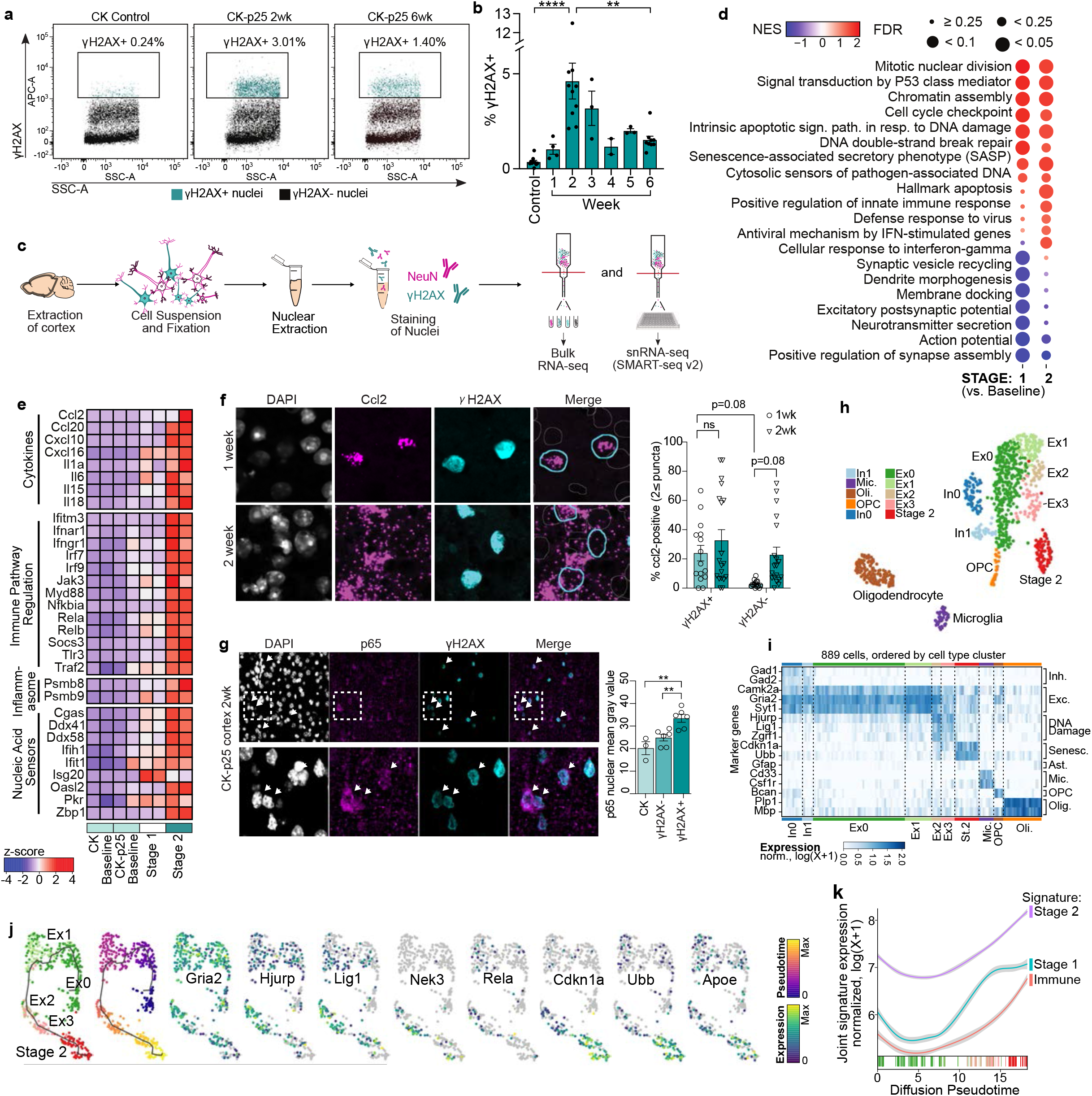
Neurons marked by DNA DSBs activate inflammatory signaling at early stages of neurodegeneration. **a.** Flow cytometry dot plots of γH2AX+ nuclei from CK and CK-p25 cortex. γH2AX+ are highlighted in turquoise, and percent total population is indicated above the gating box. **b.** CK-p25 mice were taken off dox for 1 through 6 weeks, then cortical nuclei were extracted and stained for γH2AX. Each datapoint represents percent γH2AX+ nuclei from one mouse cortex. **c.** RNA seq workflow for the gated populations. Bulk RNA-seq: Cortices were extracted from CK and CK-p25 mice (n=2 per genotype for bulk RNA-seq, 2-week timepoint only). Tissue was homogenized, then nuclei were extracted and stained for NeuN and γH2AX. 50,000 nuclei were sorted from each gated population. snRNA-seq: Cortices were extracted from CK and CK-p25 mice (n=3 per genotype×timepoint for SMART-seq, 1-week and 2-week timepoints). Tissue was homogenized, then nuclei were extracted and stained for NeuN and γH2AX. 48 nuclei were sorted for each gated CK-p25 population per mouse, and 32 nuclei were sorted for each gated CK population per mouse. **d.** Differential gene ontology terms from gene set enrichment analysis (GSEA) of bulk RNAseq data. Left column depicts GSEA results from Stage 1 vs. CK-p25 Baseline contrast. Right column depicts GSEA results from Stage 2 vs. CK-p25 Baseline contrast. Color indicates normalized enrichment score (NES). Size indicates false discovery rate (FDR). **e.** Heatmap of differentially expressed genes belonging to inflammatory gene sets from bulk RNA-seq data. Each column represents one mouse. Gene sets are organized by biological function. **f.** (left): Representative images of Ccl2 RNAscope combined with γH2AX immunofluorescent staining. Imaging was performed on 1wk and 2wk CK-p25 cortex. (right): Quantification of number of γH2AX+ and γH2AX- cells with 2≤ Ccl2 puncta. Each datapoint respresents average %Ccl2-positive cells in one image from one mouse. 4-3 images were taken per mouse. (Ccl2 1wk n=4, Ccl2 2wk n=5) **g.** Representative image of p65 immunostaining in CK-p25 cortex at two-week timepoint. White arrowheads indicate γH2AX+ nuclei with nuclear p65 immunoreactivity. Quantification of p65 mean intensity for γH2AX+ and γH2AX- nuclei. Each data point represents the average p65 nuclear mean gray value of 20-60 nuclei from one mouse. **h.** UMAP of gated populations from CK and CK-p25 cortex at 1-week and 2-week timepoints. Colors indicate cell type annotation. **i.** Marker gene expression for each cell type cluster. Columns represent 985 individual cells ordered by cell type cluster. **j.** Trajectory analysis of Ex0,1,2,3, and Stage 2 neurons. Cells from each cluster (indicated by color annotation) are ordered across pseudotime (indicated by color gradient). **k.** Trajectory analysis of Ex0,1,2,3, and Stage 2 neurons. Smoothened gene signature expression across pseudotime. Stage 1=Significantly upregulated genes from Stage 1 vs. Baseline contrast. Stage 2= Significantly upregulated genes from Stage 2 vs. Baseline contrast. Immune= Genes belonging to immune gene ontologies that were significantly upregulated in Stage 2 neurons. padj<0.05, Log2 fold-change ≥1.0. Error bars represent standard error of mean (S.E.M.); ****P<0.0001, ***P<0.001, **P<0.01, *P<0.05; One-way ANOVA with Tukey’s test for multiple comparisons (b,f). Two-way ANOVA followed by Sidak’s test for multiple comparisons (g). Data are pooled from 4 independent experiments (b). Data are representative of two independent experiments (f,g).

We found that all γH2AX-positive nuclei stained positively for CamkIIa, a marker of excitatory neurons (Supplementary Figure 1e,f). Interestingly, at the two week time point, only 57.37 ± 8.822 % of γH2AX-positive nuclei also stained positive for NeuN, a general marker for forebrain neurons (Supplementary Figure 1e,g). Degradation of neuron cell type identity is observed within aging and Alzheimer’s disease patients^31,32^, and loss of NeuN expression has been proposed to be an indicator of declining neuronal health^33^. Indeed, reduced NeuN expression or reduced NeuN immunoreactivity is an established feature of neuronal damage^34–37^. Furthermore, γH2AX-positive nuclei did not overlap with markers of microglia (Iba1), astrocytes (GFAP), or oligodendrocytes and oligodendrocyte precursor cells (Olig2), indicating that they were not of glial origin (Supplementary Figure 1h,i). Thus, we classified nuclei based on their NeuN and γH2AX immunoreactivity. This resulted in four distinct populations: γH2AX-negative NeuN-positive (“Baseline” neurons), γH2AX-positive NeuN-positive (Stage 1 neurons), γH2AX-positive NeuN-negative (Stage 2 neurons), and γH2AX-negative NeuN-negative (putative non-neuronal cells, which we refer to as “Other”) (Supplementary Figure 1g). We used fluorescence activated nuclei sorting (FANS) followed by bulk RNA sequencing to transcriptionally profile each population (Figure 1c). All nuclei were collected for sequencing at the two-week time point, which was when we observed the peak density of γH2AX-positive nuclei.

Both Stage 1 and Stage 2 populations expressed more p25 transgene compared to other populations (Supplementary Figure 1j). CK-p25 mice exclusively express p25 in excitatory forebrain neurons, as it relies on the expression of tTA driven by the CamkII promoter. Because the p25 transgene is fused to GFP, we performed staining to validate this finding. We found that 85.74 ± 0.055 % of γH2AX-positive cells, regardless of NeuN immunoreactivity, were also GFP-positive (Supplementary Figure 1k,l). To verify Stage 2 neuronal identity independent of transgene expression, we also stained for Neurod1, a neuronal transcription factor. We found that 81.96 ± 0.046 % of γH2AX-positive cells stained positively for Neurod1, regardless of NeuN immunoreactivity (Supplementary Figure 1m,n). Together, our neuron marker and transgene expression analysis indicated that both Stage 1 and Stage 2 populations were likely excitatory neurons engaged in a DSB response.

### Differential gene expression in DSB-bearing neurons

We performed two differential expression analyses to characterize gene expression changes in these populations: Stage 1 vs. CK-p25 Baseline, and Stage 2 vs. CK-p25 Baseline. Large transcriptional changes were observed in both contrasts. 3,031 upregulated and 717 downregulated transcripts were identified in Stage 1, and 5,055 upregulated and 3,792 downregulated transcripts were identified in Stage 2 (log_2_ fold change ≥ 1, adjusted p-value < 0.05) (Supplementary Figure 2a-c). Gene set enrichment analysis (GSEA)^38^ revealed that both Stage 1 and Stage 2 neurons displayed a significant enrichment for genes implicated in DSB repair, apoptotic signaling, and cell cycle, and a significant reduction in synaptic function processes (Figure 1d). These pathways were previously found deregulated in both the CK-p25 mouse and AD human brain tissue_9,26,29,39,40_.

Remarkably, we also found that a number of innate immune pathways were enriched in Stage 1 and Stage 2 neurons. This included gene ontology terms ‘Senescence-Associated Secretory Phenotype (SASP)’, ‘cytosolic sensors of pathogen-associated DNA’, and ‘positive regulation of innate immune response’ (Figure 1d). Upon closer inspection, we observed an enrichment of genes linked to DSB-mediated activation of SASP signaling, particularly in Stage 2 neurons. This included nucleic acid sensors such as *Cgas* and *Zbp1*, NFkB subcomponents *Rela* and *Relb*, SASP factors *Il6*, *Il15, Ccl2, Cxcl10,* and interferon stimulated genes *Isg15 and Ifitm3* (Figure 1e). Notably, a number of these genes are expressed in neurons following viral infection^23,24^. These genes are also classic hallmarks of senescence, suggesting that DSB accumulation elicits senescent and anti-viral-like responses in neurons.

Because the CK-p25 model is characterized by the development of type-I interferon-reactive microglia specifically by two weeks induction^26^, we wanted to determine if DSB-bearing neurons express cytokines before microglia. To do this, we performed RNAscope multiplexed fluorescent in-situ hybridization in the CK-p25 cortex after one and two weeks of induction. We focused our profiling on *Cxcl10* and *Ccl2* because these pro-inflammatory chemotactic molecules were highly expressed in Stage 2 neurons and are known to be secreted by neurons upon viral infection^23,41–44^. We also performed RNAscope for the excitatory neuron marker *Camk2a* to again confirm if γH2AX-positive nuclei were of neuronal origin. In agreeance with our RNAseq and flow cytometry data, 98.81 ± 1.19% of γH2AX-positive nuclei were also *Camk2a*-positive (Supplementary Figure 2d). This high positivity rate reveals that γH2AX-positive nuclei are nearly exclusively neurons. RNAscope analysis also revealed that DSB-bearing neurons were the only cells to express *Cxcl10* and *Ccl2* at the one-week time point. In contrast, *Cxcl10* and *Ccl2* gene expression was highly enriched in other cell types at the two-week time point (Figure 1f, Supplementary Figure 2e). This indicated that expression of these chemokines in γH2AX-positive neurons precedes their expression in glial cells, and that cytokine secretion from DSB-bearing neurons may be an early mechanism of glial cell recruitment and activation in the CK-p25 brain.

To identify the master regulators of DSB-associated neuronal immune signaling, we performed transcription factor enrichment analysis using Enrichr^45–47^. A subset of immune pathway genes was extracted from the significantly upregulated genes in Stage 2 neurons for analysis. The overlap between this immune gene module and transcription factor target genes was then calculated. Multiple subunits of the NFkB complex were consistently enriched across Enrichr transcription factor libraries, including Rela, Nfkb1, and Relb (Supplementary Table 1). Notably, the NFkB transcription factor plays a well-established role in SASP activation and the DSB response^19,20,22^. We chose to focus on Rela, also known as p65, because it is a core member of the canonical NFkB complex, and it was most frequently enriched in the Enrichr analysis. We stained for p65 in two-week CK and CK-p25 cortices. NFkB is normally sequestered in the cytosol, but translocates to the nucleus to form an active complex upon cellular insult. Nuclear p65 intensity was significantly higher in γH2AX-postive neurons compared to other γH2AX-negative cells (Figure 1g), supporting evidence of increased NFkB transcriptional activity.

### Single cell RNA-sequencing if DSB-bearing neurons

Notably, inflammatory gene expression was lower in Stage 1 neurons compared to Stage 2 neurons (Figure 1e), and more Stage 1 neurons were identified at the one-week time point compared to Stage 2 neurons (Supplementary Figure 2f,g). This suggested that the Stage 2 population may develop after Stage 1, which aligned with our initial prediction when defining these distinct cellular states based on NeuN-reactivity. To better understand the relationship between Stage 1 and Stage 2 neurons, we performed single nucleus RNA-sequencing on each FANS-gated population at both one and two-week timepoints (Figure 1c). A total of 1,357 single nucleus libraries were prepared using SMARTseq2 chemistry. Following quality control measures (see methods), 889 libraries remained for downstream analysis (Figure 1h, Supplementary Figure 3a). Cells were classified into major cell type clusters based on marker gene expression, resulting in the identification of 521 excitatory neurons, 71 inhibitory neurons, 131 oligodendrocytes, 33 oligodendrocyte precursor cells (OPCs), and 50 microglia. We did not detect any astrocytes.

Remarkably, the majority of Stage 2-gated nuclei formed their own cell type cluster (Figure 1h). While we did identify 13 Stage 2-gated nuclei in microglia and oligodendrocyte clusters, all of the nuclei in question came from one CK-p25 mouse (Supplementary Figure 3b,c). None of the other five CK-p25 mice had Stage 2-gated nuclei fall into glial clusters. Combined with our previous cell type immunostaining analysis (Supplementary Figure 1h,i), we concluded that the immune signature identified in the Stage 2 population was not likely to be driven by contamination from microglia or oligodendrocytes. Compared to other neuronal clusters, the *in silico* Stage 2 cluster was significantly enriched for the FANS-gated Stage 1 and Stage 2 gene signatures (Supplementary Figure 3d). Interestingly, the Stage 2 cluster expressed only moderate levels of the excitatory neuron cluster markers *Camk2a*, *Gria1*, and *Syt2*, and lacked marker genes of other canonical cell types such as inhibitory neurons (*Gad1*, *Gad2*), astrocytes (*GFAP*), microglia (*Cd33*, *Csf1r*), oligodendrocytes (*Plp1*, *Mbp*), and OPCs (*Bcan*). Instead, the Stage 2 cluster expressed marker genes indicative of senescence, including *Cdkn1a,* and *Ubb* (Figure 1i). These distinctive cell type markers further suggested to us that the Stage 2 cluster represented a population of DSB-bearing neurons engaged in a senescence-like inflammatory response.

To further examine Stage 1 and Stage 2 cell type heterogeneity, we subclustered all neuronal cells. This resulted in the identification of four excitatory neuron subclusters (Ex0-3), and two inhibitory neuron subclusters (In0, In1) (Figure 1h). Stage 1 cells were enriched in subclusters Ex2 and Ex3 compared to Ex0 and Ex1 (Supplementary Figure 3c). Subclusters Ex2 and Ex3 were also marked by DNA repair-associated genes, such as *Hjurp*, *Lig1*, and *Zgrf1*, further suggesting that Ex2 and Ex3 cells were engaged in a DSB repair response (Figure 1i).

Interestingly, these DNA repair-associated genes were not expressed in the Stage 2 cluster. Combined with the enrichment for the classical senescence marker *Cdkn1a*, this suggested that Stage 2 cells may be engaged in a later-stage response to DSBs. This motivated us to perform trajectory analysis. The single cell analysis package Monocle3^48–50^ was used to order cells from subclusters Ex0,1,2,3 and Stage 2 along a pseudotime, which was generated through a learned trajectory of subcluster gene expression differences (Figure 1j). Both CK and CK-p25 cells were used for this analysis. Next, we identified genes that changed as a function of pseudotime. We observed sequential enrichment of Stage 1 and Stage 2 gene signatures along pseudotime (Figure 1k). The expression of Stage 2 signature genes specifically involved in immune response (Immune) were also enriched at the end of pseudotime (Figure 1k). We found that genes associated with neuronal identity such as *Gria2* and *Kalrn* were most highly expressed at the beginning of the trajectory (Figure 1j). Meanwhile, genes associated with DNA repair like *Hjurp*, *Lig1*, and *Nek3* peaked along the middle, and *Cdkn1a* expression peaked at the end of the trajectory. We also observed late-stage expression of *Apoe*, supporting previous observations that *Apoe* expression increases in neurons following injury^51^.

### DSBs elicit innate immune signaling in neurons

To test if this late-stage DSB response signature is triggered by DSBs or represents a unique response mounted by CK-p25 mice, we treated murine primary neuron cultures with 50uM etoposide (ETP). ETP produces DSBs by inhibiting topoisomerase-II function, thus preventing the re-ligation of cleaved DNA-TopII complexes (Figure 2a). This dose of ETP was consistent with previous studies that used ETP to investigate DSB-elicted immune activity *in vitro*^52^. We also performed cell type immunostaining analysis to assess the cell type composition of our primary cultures. 82.87% of cells were identified as mature neurons (NeuN-positive), and another 9.31% were neural precursor cells (Nestin-positive), (Supplementary Figure 4a). A trace number of cells were also identified as astrocytes (GFAP-positive, 2.46%) and oligodendroglia (Olig2-positive Nestin-negative, 1.35%).

**Figure 2.**
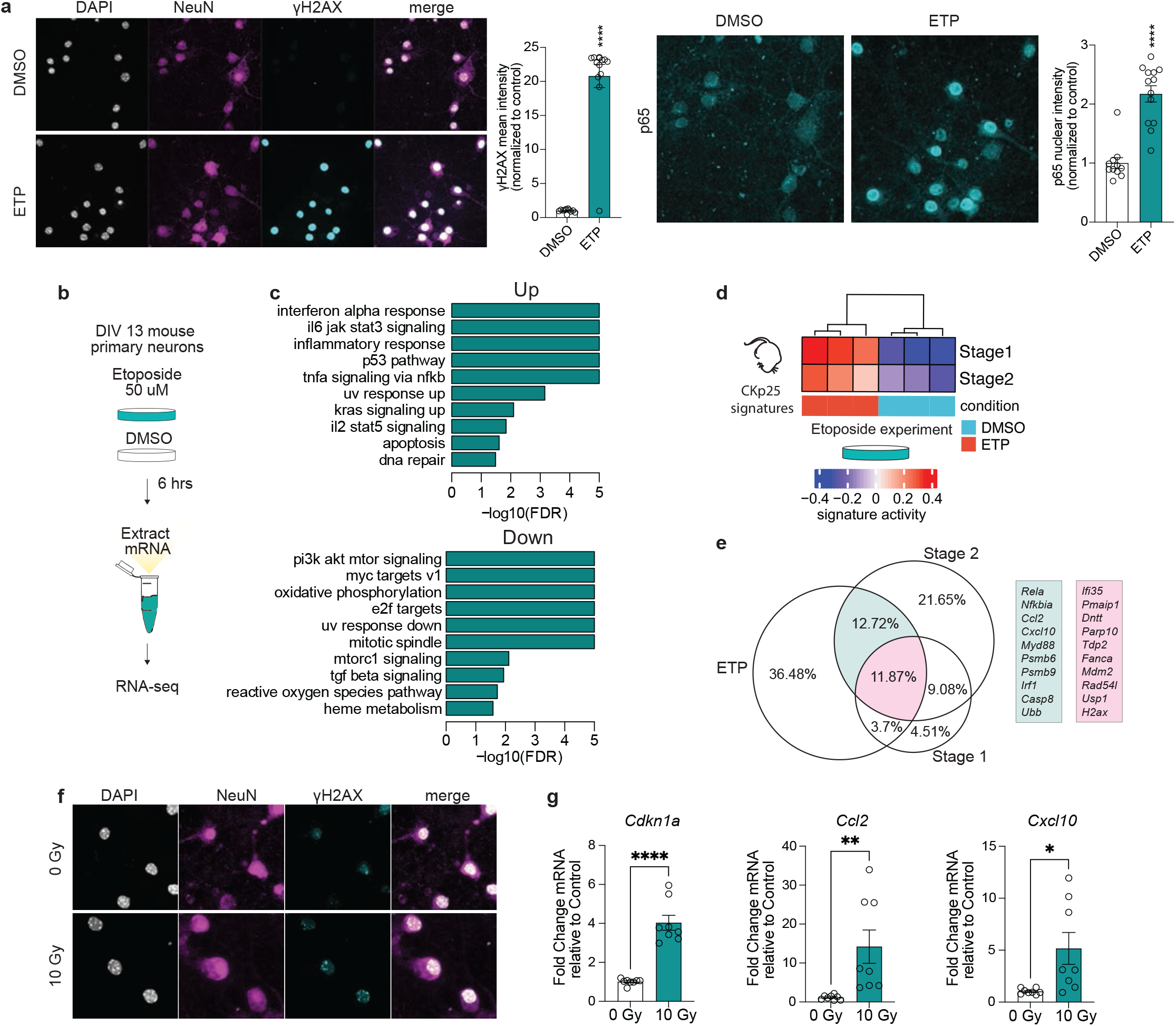
Induction of DNA DSBs is sufficient to elicit immune pathway signaling in neuron primary culture. **a.** Top: Representative images of NeuN and γH2AX immunostaining in ETP and vehicle-treated primary cultures. γH2AX immunoreactivity is quantified for each condition. Each data point represents γH2AX mean gray value for one nucleus in the representative image. Bottom: Representative images of p65 immunostaining in ETP and vehicle-treated primary cultures. p65 immunoreactivity is quantified for each condition. Each data point represents p65 mean gray value for one nucleus in the representative image. **b.** Schematic of etoposide (ETP) treatment. DIV13 neuron primary culture were treated with either 50uM ETP or vehicle control (DMSO) for 6 hours. mRNA was extracted from the cultures and sequenced. **c.** Differential gene ontology terms identified through gene set enrichment analysis (GSEA) from ETP vs. DMSO contrast. **d.** Heatmap of Stage 1 and Stage 2 signature enrichment (from bulk RNAseq) in DMSO and ETP-treated neurons. **e.** Venn diagram of significantly upregulated protein-coding genes from ETP-treated neurons and Stage 1 and Stage 2 gene signatures. Percentages are in reference to the total number of unique genes from all three gene sets. Example genes are shown to the right. Genes overlapping in ETP and Stage 2 are in turquoise. Genes overlapping in ETP, Stage 2, and Stage 1 are in magenta. The area of the circles are in proportion to the size of the gene sets. **f.** Representative images of NeuN and γH2AX immunostaining from 10 Gy and 0 Gy x-ray irradiation treated primary neurons. **g.** qRT-PCR of Cdkn1a, Ccl2, and Cxcl10 in 10 Gy irradiated primary neurons. Each datapoint represents one biological replicate. Error bars represent standard error of mean (S.E.M.); ****P<0.0001, ***P<0.001. Student’s T-test (a,g). Data are representative of 3 independent experiments (a). Data are representative of 2 independent experiments (g).

RNA sequencing was performed to profile the transcriptional changes occurring in primary neurons after ETP treatment (Figure 2b). There was a significant upregulation of 7,532 transcripts and downregulation of 7,106 transcripts compared to vehicle-treated control neurons (DMSO) (adjusted p-value<0.05) (Supplementary Figure 4b-d). Notably, GSEA revealed that biological pathways mediating senescent and antiviral activity such as ‘interferon alpha response,’ and ‘il6 jak stat3 signaling,’ were enriched in ETP-treated cultures (Figure 2c). Furthermore, many of the cytokines upregulated in Stage 2 neurons were also upregulated in ETP-treated neurons, including *Ccl2*, *Cxcl10*, *Cxcl16*, and *Il6* (Supplementary Table 2). Stage 1 and Stage 2 gene signatures were both significantly enriched (p=0.00064 & p=0.0081 respectively) (Figure 2d, Supplementary Figure 4e). To perform a more detailed comparison of Stage 1 & 2 signature genes to ETP DEGs, we generated a Venn-Diagram (Figure 2e). 11.87% of all DEGs (total DEGs from all datasets) were shared between ETP, Stage 1 and Stage 2 conditions. This overlap was enriched for DNA damage repair genes such as *Parp10*, *Tdp2*, *Fanca*, and *Rad54l*. A further 12.87% of all DEGs were shared between ETP and Stage 2 conditions. This overlap was enriched for immune genes such as *Rela*, *Nfkbia*, and *Irf1*.

To confirm immune genes were expressed by neurons and not by other trace cell types in our primary cultures, we performed RNAscope for *Ccl2* and the gene encoding NeuN, *Rbfox3*. Cells that were *Rbfox3*-positive had significantly more *Ccl2* puncta than *Rbfox3*-negative cells (Supplementary Figure 4f). p65 nuclear intensity was also significantly increased in ETP-treated neurons (Figure 2a). A NeuN co-stain was used to confirm p65 activation was derived from neurons.

We also used an independent method of DSB induction in neurons by treating primary cultures with 10 Gy of X-ray irradiation (Figure 2f). After a 24-hour recovery period, we performed RT-qPCR to assess the mRNA levels of the Stage 2 marker gene *Cdkn1a*, and the antiviral signaling genes *Ccl2* and *Cxcl10*. 10 Gy of X-ray irradiation significantly increased the expression of all three of these genes (Figure 2g).

### A DSB-associated immune signature is conserved in human neurons

We next sought to determine if signatures of DSB-bearing neurons can also be detected in the human brain. First, we used the Stage 2 gene signature to identify neurons with DSB-associated immune activation in a previously published snRNA-seq dataset of individuals with and without AD pathology (Figure 3a)^53^. We assessed Stage 2 gene expression in all major brain cell types: excitatory neurons, inhibitory neurons, astrocytes, oligodendrocytes, oligodendrocyte precursor cells, and microglia. For each cell type, genes were ranked based on their expression and correlation to ‘global pathology,’ a variable quantifying three major AD pathologies: neuritic plaques, diffuse plaques, and neurofibrillary tangles^54^. Genes positively correlated with global pathology for each cell type were then sampled for the enrichment of Stage 2 genes (Figure 4b). The strongest enrichment of Stage 2 genes was identified in excitatory neurons (p=8.7E-19), and inhibitory neurons (p=7.7E-7). Astrocytes and oligodendrocytes were also found to have a significant enrichment of Stage 2 genes correlating with global pathology, but to a much lesser extent (Figure 4b), (Bonferroni adjusted p-value < 0.01). Therefore, while we cannot exclude the possibility that human astrocytes and oligodendrocytes may carry some disease-associated enrichment for Stage 2 genes, the Stage 2 signature is largely driven by neurons in this dataset. When assessing the Stage 2 genes positively correlated with global pathology in excitatory neurons, we again observed an enrichment of immune-related biological pathways, including the genes Ccl2-like receptor (*CCRL2*), *CD74*, *CXCL16*, and *APOE* (Figure 4c,d).

**Figure 3.**
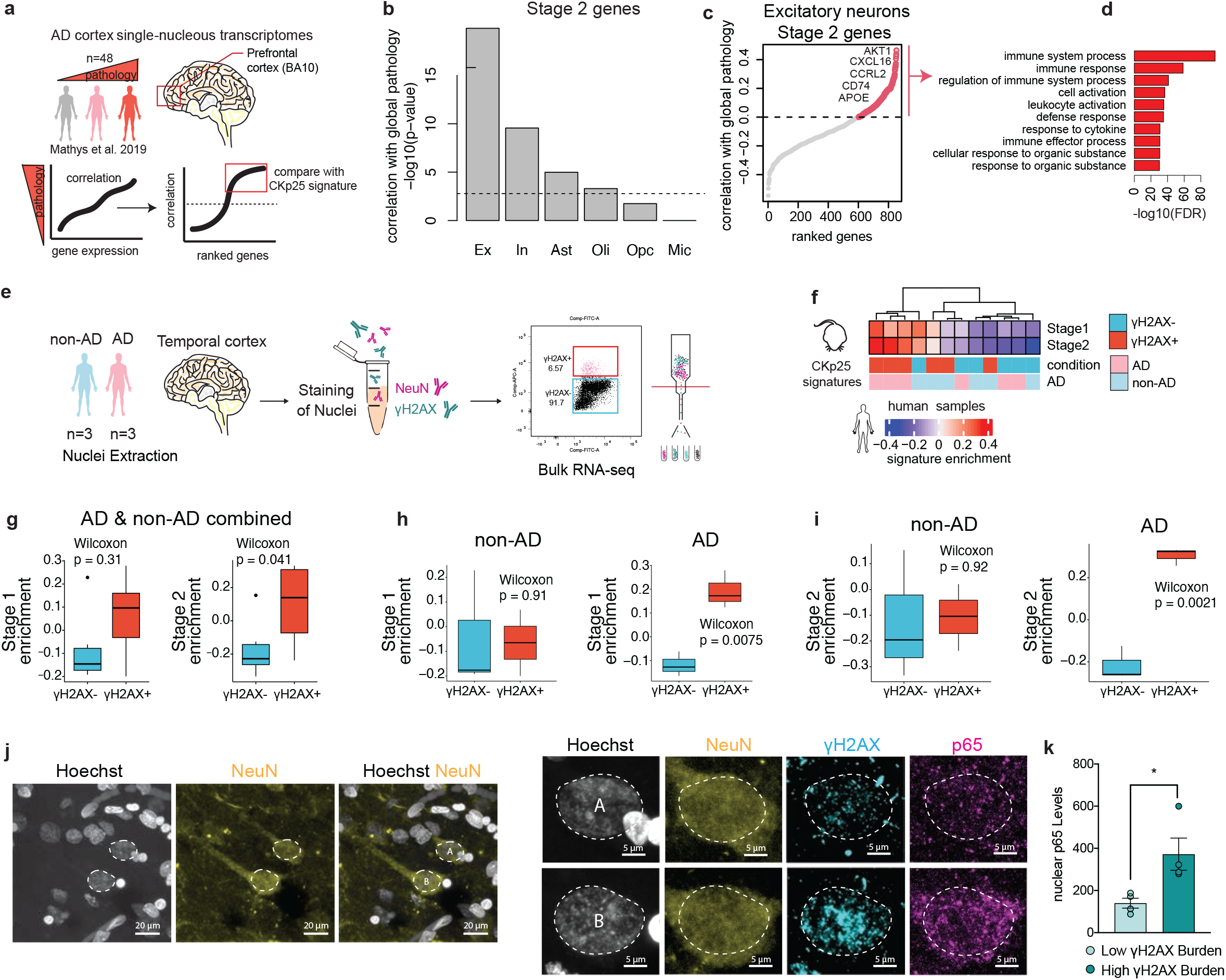
Inflammatory signaling in DSB-bearing neurons is positively correlated with Alzheimer’s disease pathology. **a.** Schematic of the Stage 2 signature analysis in the snRNA-seq dataset from Mathys et al., 2019. **b.** Quantification of Stage 2 signature correlation with global pathology in celltype clusters from Mathys et al., 2019. It was tested if stage 2 genes were significantly and positively correlated with the global pathology metric for each major celltype. The -log10 p-value for these tests are shown in the histogram. The dashed line indicates a p-value of 0.01 after Bonferroni correction for multiple testing. Excitatory neurons (Ex), Inhibitory neurons (In), Astrocytes (Ast), Oligodendrocytes (Oli), Oligodendrocyte precursor cells (Opc), Microglia (Mic). **c.** Stage 2 signature genes ranked by their correlation with global pathology in excitatory neurons. Stage 2 genes with positive correlation are shown with red circles. **d.** Gene ontology enrichment of Stage 2 signature genes positively correlated with global pathology. **e.** Schematic of γH2AX+ nuclei sorting from AD and non-AD brain tissue. **f.** Heatmap of Stage 1 and Stage 2 signature enrichment in γH2AX+ and γH2AX- human NeuN+ nuclei. **g-i.** Quantification of Stage 1 and Stage 2 signature enrichment in γH2AX+ and γH2AX- human NeuN+ nuclei samples by FANS gate (g), and FANS gate and disease status (h,i). **j.** Representative image of γH2AX, p65, and NeuN in the AD brain. Left: two NeuN-positive nuclei are outlined (white dashed line). Right: magnification of the two outlined nuclei. Top nucleus (A) represents low γH2AX burden, bottom nucleus (B) represents high γH2AX burden. **k.** Quantification of p65 nuclear enrichment in low and high γH2AX-burdened neurons. Each dot represents the average of 23-41 NeuN+ nuclei per individual. Error bars represent standard error of mean (S.E.M.); ****P<0.0001, ***P<0.001, **P<0.01, *P<0.05, ns not significant. T-test (k). Wilcoxon test (g-i).

**Figure 4.**
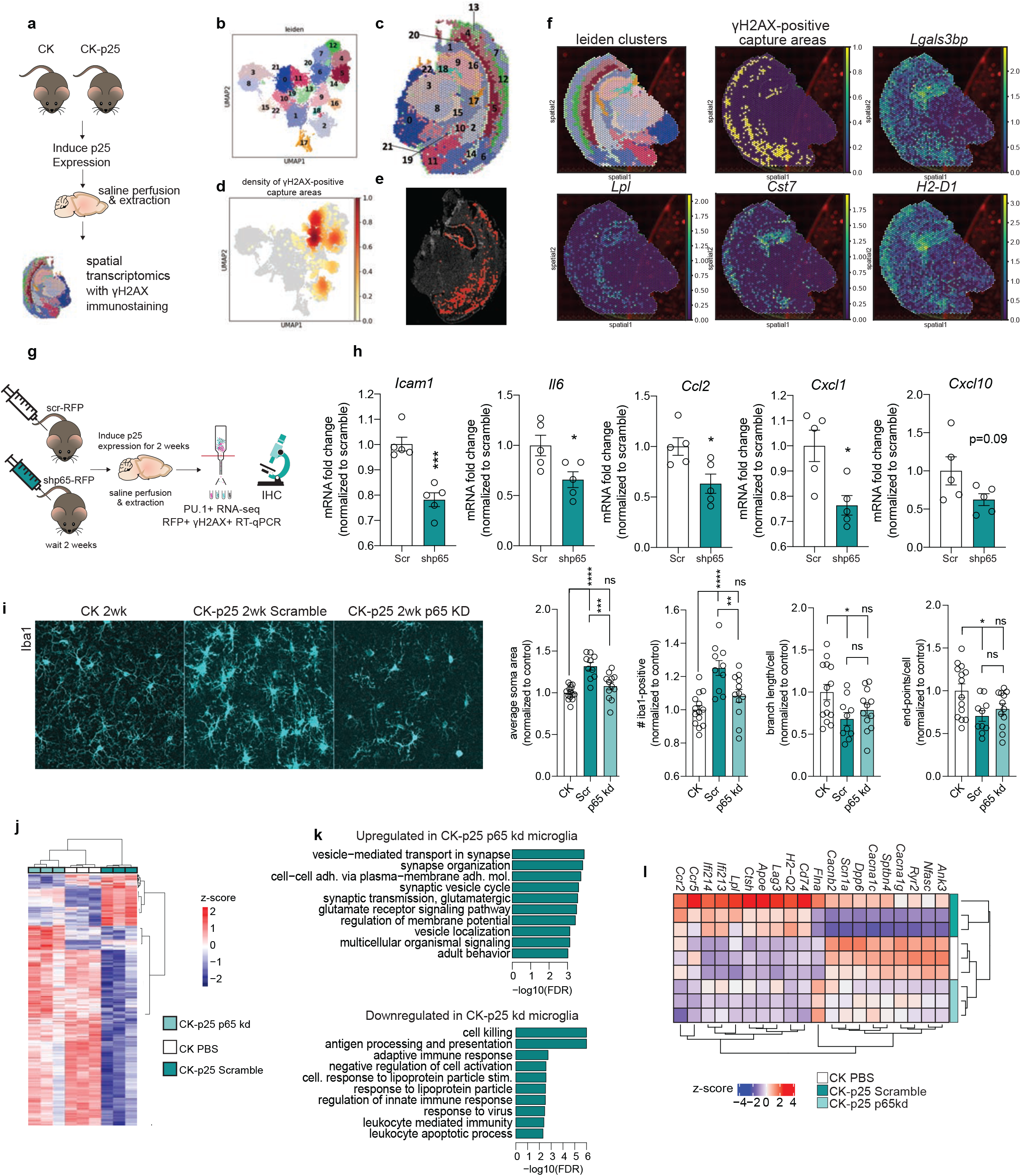
Immune signaling in neurons recruits and activates microglia. **a.** Schematic of spatial transcriptomics experiment. CK (n=3) and CK-p25 (n=4) were induced for two weeks. Coronal brain sections were stained and imaged for γH2AX, then sequenced. **b.** UMAP of capture areas from all samples. Leiden clusters are indicated by color and number. Each dot represents one capture area. **c.** Leiden clusters superimposed onto a CK-p25 brain slice used for spatial transcriptomics. **d.** UMAP indicating the density of γH2AX-positive capture areas. **e.** γH2AX-positive capture areas identified in one CK-p25 brain slice. **f.** Spatial clusters, γH2AX-positive capture areas, and reactive microglia signature gene expression in one CK-p25 sample. **g.** Schematic of neuronal p65 knock-down experiment. CK-p25 mice receive retro-orbital injections of scramble shRNA-RFP AAV or shp65-RFP AAV. CK mice received retro-orbital injections of PBS. Mice were allowed to recover for two weeks before being taken off dox. Brains were collected for subsequent analysis at the two week timepoint. **h.** qRT-PCR of immune genes in sorted γH2AX+ nuclei. **i.** Representative images of Iba1 immunostaining in CK, Scramble, and p65kd cortex. **j.** Quantification of (left to right): Iba1+ soma area, number Iba1+ per image, average branch length per Iba1+ cell, and number end-points per Iba1+ cell. Each data point represents one image. Two images were taken per mouse. **k.** Heat map of differentially expressed genes from P65 vs. Scramble contrast. Each column represents one mouse. **l.** Upregulated and downregulated gene ontology (biological pathway) terms identified through gene set enrichment analysis (GSEA) from p65 kd vs. scramble contrast. **m.** Heatmap of significantly upregulated and downregulated genes in Pu.1+ nuclei from p65 kd cortex compared to Pu.1+ nuclei from scramble cortex. Each row represents gene expression from one mouse. Error bars represent standard error of mean (S.E.M.); ****P<0.0001, ***P<0.001, **P<0.01, *P<0.05; n.s. not significant. Student’s t-test (h). One-way ANOVA followed by Holm-Sidak’s test for multiple comparisons (i). h: Scramble (n=5), p65kd (n=5). i: CK (n=7), Scramble (n=5), p65kd (n=6).

To validate the enrichment of DSB-bearing neurons identified in our snRNA-seq analysis, we sorted γH2AX-positive and γH2AX-negative NeuN-positive and NeuN-negative nuclei from the postmortem temporal cortex of 6 individuals, and then performed bulk RNA sequencing (Figure 4e, Supplementary Figure 5a-c). These individuals were part of the Massachusetts Alzheimer’s Disease Research Center (MADRC) cohort, and were divided by Braak score as either AD (Braak 4-6), or non-AD (Braak 1-3). Age, sex, and other biological variables for each individual are available in Supplementary Table 3. To assess the cell type specificity of our sorted populations, we examined the expression of several cell type marker genes (Supplementary Figure 5d-f). Unlike the Stage 2 nuclei from the CK-p25 mice, γH2AX-positive NeuN-negative nuclei from the human brain were enriched for glial markers (Supplementary Figure 5e,f). This indicated to us that human glial cells also accumulate DSBs, and that bulk γH2AX-positive NeuN-negative nuclei were likely to contain a heterogenous population of many different cell types. Thus, to ensure we were analyzing neuronal gene expression, we focused our analysis on γH2AX-positive NeuN-positive nuclei.

We assessed Stage 1 and Stage 2 gene signature activity in NeuN-positive nuclei from AD and non-AD human brains (Figure 4f). γH2AX-positive neurons displayed modest enrichment for the CK-p25 Stage 1 and Stage 2 genes (p=0.31 and p=0.04 respectively) (Figure 4g). Comparing samples by AD status and γH2AX status further revealed that Stage 1 and Stage 2 gene activity was markedly enriched in γH2AX-positive neurons from AD individuals (p=0.0075 and p=0.0021, respectively) (Figure 4h,i). Furthermore, when contrasting all γH2AX-positive neurons against all γH2AX-negative neurons, we found that the average log_2_ fold change values for both Stage 1 and Stage 2 gene sets were positive, and significantly higher than the average log_2_ fold change values for all other genes (Supplementary Figure 5g). We also assessed the transcriptional similarity between human γH2AX-positive neurons and ETP-treated primary neurons. Again, we found striking similarities between the DSB signature produced in the human brain, and the DSB signature generated by ETP treatment of primary neurons (Supplementary Figure 5h).

To further verify the enrichment of immune activation in DSB-bearing neurons from brains with AD pathology, we used another cohort of AD brain tissue to assess nuclear p65 expression in NeuN-positive neurons with high versus low burdens of DSBs. We acquired 4 prefrontal cortex tissue samples from individuals belonging to the Religious Orders Study cohort. All individuals analyzed exhibited AD pathology, described in Supplementary Table 4. To bin neurons by DSB burden, we analyzed γH2AX expression in NeuN-positive nuclei. The median γH2AX expression was calculated for each brain, and used to split neurons into the high or low DSB condition (Supplementary Figure 6a). Neurons with high DSB burden consistently expressed higher levels of nuclear p65 compared to neurons with low DSB burden (Figure 3j,k, Supplementary Figure 6b).

In summary, through the analysis of an independent snRNA-seq dataset, FANS RNA-sequencing, and immunohistochemistry, we found that SASP and antiviral-like pathways were active in human DSB-bearing neurons. Furthermore, this neuronal immune signature was further amplified in the context of AD pathology, suggesting that it may serve a functional role in disease-associated neuroinflammation.

### DSB-bearing neurons stimulate glial activation

We have previously published an in-depth characterization of reactive microglia in CK-p25 mice using snRNA-seq^26^. We wondered if these microglia were responding to immune signaling from DSB-bearing neurons. To address this, we performed γH2AX immunostaining followed by spatial transcriptomics on two-week induced CK-p25 brains (Figure 5a). The capture areas of 10X Visium spatial transcriptomics slides are large enough to encompass the transcriptional profile of 3-5 cells in mouse brain tissue. Therefore, we hypothesized that capture areas with γH2AX signal would also contain transcripts from nearby microglia. First, we identified 23 spatial clusters across three CK and four CK-p25 coronal sections (Figure 4b,c, Supplementary Figure 7a,b). Next, we used a binary classification system to identify capture areas with γH2AX signal past a given threshold (See Methods, Figure 4d,e, Supplementary Figure 7c). To determine if the reactive microglia characterized in Mathys et al., 2017 were enriched in γH2AX-positive capture areas, we performed a differential comparison between all γH2AX-positive capture areas and all γH2AX-negative capture areas. Using GSEA, we identified a robust enrichment of the microglia signature (1,383 significantly upregulated genes in reactive microglia compared to homeostatic microglia) within γH2AX-positive capture areas compared to γH2AX-negative capture areas (Supplementary Figure 8a). This indicated that overall, the reactive microglia signature characterized in Mathys et al., 2017 was significantly associated with γH2AX-positive neurons.

**Figure 5.**
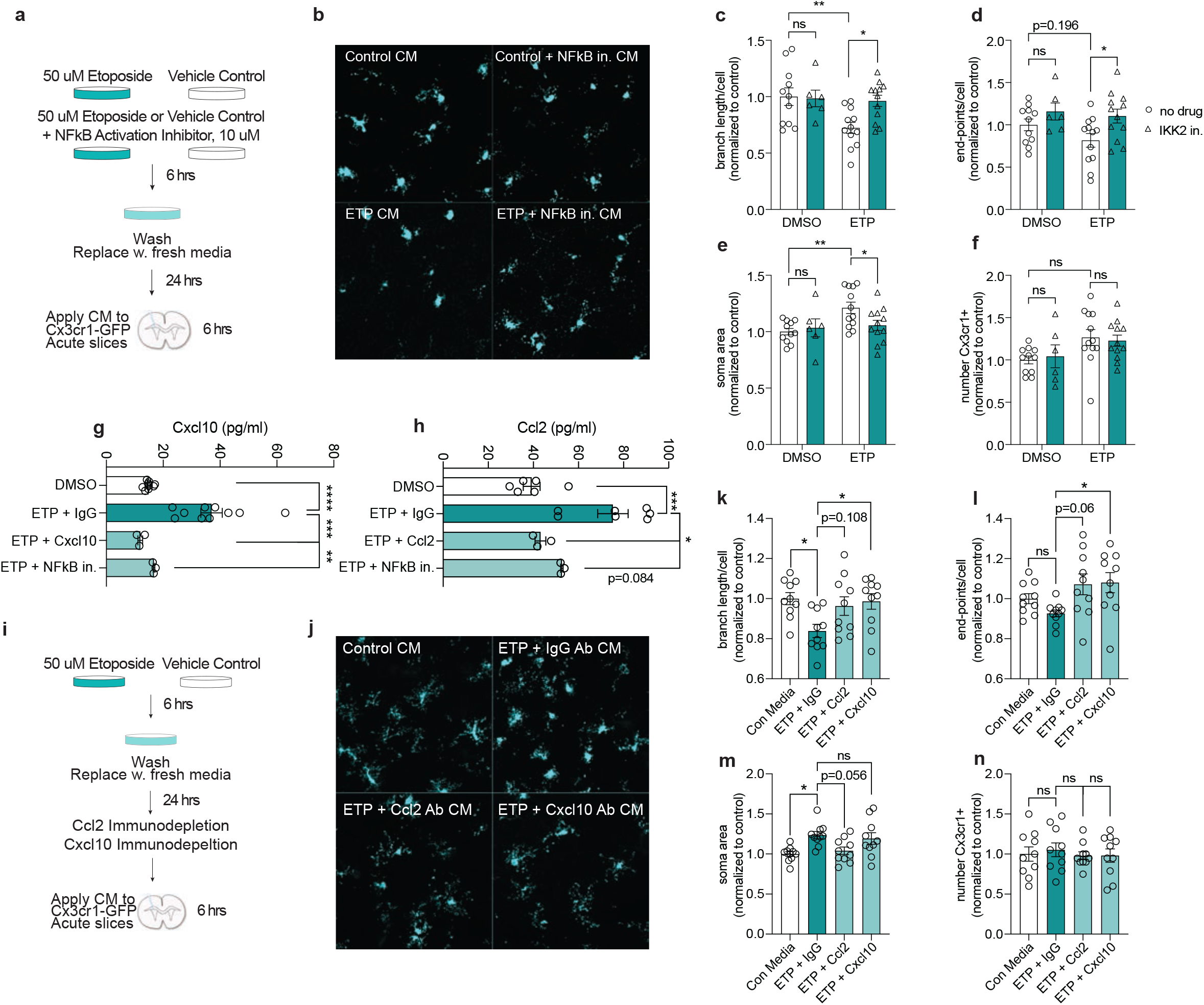
Ccl2 and Cxcl10 are secreted from DSB-bearing neurons to activate microglia. **a.** Schematic for treating acute Cx3cr1-GFP slices with conditioned media from etoposide-treated primary neuron cultures. Cultures were either treated with 50uM ETP or vehicle control (DMSO) for six hours. In a separate condition, primary neurons were treated with 50uM ETP and 10uM NF-kappaB Activation Inhibitor VI (IKK2 inhibitor). Cultures were washed with PBS after six hours and media was replaced. After 24 hours, this media was applied to acute Cx3cr1-GFP slices for 6 hours. **b.** Representative images of GFP in Cx3cr1-GFP acute slices treated with primary neuron conditioned media. **c-f.** Quantification of branch length per microglia (c), end-points per microglia (d), soma area (e), and number of microglia per image (f). Each data point represents the average measurement from two images in one acute slice. **g,h.** Quantification of Cxcl10 (g) and Ccl2 (h) from conditioned media from control and etoposide-treated primary neurons. Each datapoint represents one biological replicate. **i.** Schematic of ETP conditioned media experiment. Primary neurons were treated with either ETP or DMSO for 6 hours, washed with PBS, then media was replaced. Cultures recovered for 24 hours before conditioned media was collected. IgG, Ccl2, or Cxcl10 antibodies were used to immunodeplete conditioned media before they were applied to Cx3cr1-GFP acute slices for 6 hours. **j.** Representative images of microglia from acute slices treated with different conditioned media. **k-n.** Quantification of (k) branch length per microgli (l) end-points per microglia, soma area (m), and number of microglia per image (n). Each data point represents the average measurement from two images in one acute slice. Error bars represent standard error of mean (S.E.M.); ****P<0.0001, ***P<0.001, **P<0.01, *P<0.05, ns not significant. Two-way ANOVA followed by Sidak’s test for multiple comparisons (c-f). One-way ANOVA followed by Tukey’s test for multiple comparisons (k-n). Data are combined from two independent experiments (c-f, k-n). Data are combined from three independent experiments (h,i).

By mapping the density of γH2AX-positive capture areas across all spatial clusters, we further prioritized distinct spatial clusters enriched for γH2AX-positive capture areas (Figure 4d, Supplementary Figure 8b). We focused on spatial clusters with γH2AX-positive capture areas comprising 20% or more of total capture areas (Supplementary Figure 8b). This included clusters 16 and 18, which corresponded to CA1,2,3 and the dentate gyrus respectively, and clusters 2,5,6,7, which were primarily localized to the cortex. Differential comparisons of γH2AX-positive capture areas and γH2AX-negative capture areas within each cluster of interest revealed a significant enrichment of the reactive microglia signature (Supplementary Figure 8a). Notably, many of the marker genes for reactive microglia displayed expression patterns that visually correlated with γH2AX-positive capture areas, including the galectin gene *Lgals3bp*, lipase *Lpl*, cystatin *Cst7*, and MHC Class I gene *H2-D1* (Figure 4f). Genes also enriched in DSB-bearing neurons, such as *Cxcl10* and *Ifitm3* was also closely correlated with γH2AX-positive capture areas (Supplementary Figure 9). Collectively, this spatial transcriptomic analysis indicated that reactive microglia are closely associated with γH2AX-positive neurons, suggesting the occurrence of neuro-immune communication between damaged neurons and microglia.

To determine if microglia responded to DSB-mediated immune signaling, we sought to inhibit a master regulator of immune gene expression in neurons. Our data revealed that NFkB transcription factor activity is increased in DSB-bearing neurons, as are the genes downstream of NFkB activation. Therefore, we focused on inhibiting NFkB transcriptional activity. To achieve this in the CK-p25 mouse, we opted for brain-wide viral delivery of shRNAs targeting p65^55^. We performed retro-orbital injections of PHP.eb AAV shp65-RFP or Scramble-RFP into CK-p25 mice. CK control mice were injected with phosphate-buffered saline. Two weeks after injections, mice were taken off doxycycline and induced for two weeks (Figure 4g).

The majority of RFP co-localized with NeuN and γH2AX immunoreactivity, indicating neurons were the primary targets of shRNA expression, and confirming the previously reported tropism of PHP.eb for neuronal infectivity (Supplementary Figure 10a,b)^55^. The reduced expression of p65 in mice was confirmed via qPCR and immunofluorescent staining (Supplementary Figure 10c-e). The number of γH2AX-positive neurons remained the same between p65 knock-down (p65kd) and Scramble mice (Supplementary Figure 10f,g). Next, we sorted γH2AX+ and γH2AX- RFP+ neurons from both p65kd and Scramble mice to assess changes in immune gene expression (Supplementary Figure 11a). RT-qPCR analysis revealed that a number of immune genes were downregulated in DSB-bearing neurons from p65kd mice compared to Scramble, including *Ccl2, Icam1, Il6*, and *Cxcl1* (Figure 4h). Downregulation of *Ccl2* in DSB-bearing neurons was also confirmed via RNAscope (Supplementary Figure 11b,c). This indicated p65kd was able to alter neuron immune gene signatures.

To determine if suppressing neuron immune genes rescued microglial activation, we performed immunofluorescent staining with the microglia marker Iba1. We found that p65kd mice showed a reduced number of microglia (Figure 4i), which was confirmed through flow cytometry analysis of PU.1-positive nuclei from the entire cortex (Supplementary Figure 11d,e). We also found that microglia soma area was significantly reduced, but differences in branch length and end points were not statistically significant (Figure 4i). These data show that suppressing neuron immune signaling significantly affects the number and morphology of microglia.

To determine if changes in microglial number and morphology were accompanied with changes in gene expression, we performed RNA sequencing on sorted PU.1-positive nuclei from CK control, CK-p25-scramble, and CK-p25-p65kd cortex (Figure 4g, Supplementary Figure 11d). Differential analysis comparing microglia from p65kd and Scramble cortices revealed the upregulation of 627 transcripts and downregulation of 160 transcripts (Figure 4j, Supplementary Figure 11i,j). Interestingly, pathways related to synaptic activity and membrane organization were significantly upregulated in microglia from p65kd mice. This included calcium channels *Cacna1g*, *Cacna1c*, and *Cacnb2*, and sodium channel *Scn1a*, suggesting p65kd microglia may return to a more homeostatic state embodied by CK microglia (Figure 4j). In contrast, the genes that were downregulated in microglia from p65kd mice were involved in cell killing and antigen processing, such as *H2-Q2*, *Lag3*, *Ccr5*, and *Ccr2*. Notably, the *Ccr2* gene encodes the receptor for Ccl2, further implying an alteration of the Ccl2-Ccr2 axis (Figure 4k,l). We also observed a downregulation of genes related to lipoprotein processing (Figure 4k,l). Together, these results suggest that neuron immune signaling disrupts microglia homeostatic activity, and activate cell killing and antigen processing mechanisms in microglia.

Finally, to determine if specific cytokines expressed and secreted by DSB-bearing neurons play a role in microglia activation, we transitioned to our *in vitro* ETP model of DNA damage. First we wanted to understand how conditioned media from ETP-treated neurons affect microglia morphology. To do this, we generated organotypic acute brain slice cultures from Cx3cr1-GFP mice, which express GFP in all microglia. After ETP treatment and washout with PBS, neurons recovered in fresh media for 24 hours. This conditioned media was then collected and applied to Cx3cr1-GFP brain slices for 6 hours (Figure 5a). The conditioned media from ETP-treated neurons reduced microglia branch length (Figure 5b,c), but did not significantly affect the number of endpoints per microglia (Figure 5d). We also observed increased soma area (Figure 5e). Treating primary neurons with an NFkB activation inhibitor before and during ETP treatment significantly reduced the soma area of Cx3cr1-GFP-positive microglia, and increased branch length and end-points (Figure 5b-e). Conditioned media from ETP-treated neurons did not elicit a robust difference in the number of microglia analyzed per image (Figure 5f). These data further demonstrate NFkB activity as a significant mediator of immune signaling in DSB-bearing neurons.

Notably, conditioned media from ETP-treated neurons was significantly enriched for Cxcl10 and Ccl2 (Figure 5g,h). Furthermore, the NFkB activation inhibitor reduced the concentration of these cytokines (Figure 5g,h). Previously, we found that DSB-bearing neurons are the first cells to express Cxcl10 and Ccl2 in the CK-p25 cortex (Figure 1f, Supplementary Figure 2e). These cytokines induce the migration of microglia and macrophages to sites of viral infection, traumatic brain injury, and gliomas, suggesting that they may be primary constituents of immune signaling in DSB-bearing neurons^23,43,44,56,57^. To determine if they played a role in DSB-mediated microglia activation, we immunodepleted Ccl2 or Cxcl10 from conditioned media (Figure 5i). Both Ccl2 and Cxcl10 depletion increased branch length and end-points per microglia compared to IgG control depletion (Figure 5j,k,l). Interestingly, Ccl2 depletion reduced microglia soma area, but Cxcl10 depletion did not (Figure 5m). Neither immunodepletion had an effect on microglia number (Figure 5n). Combined, this suggests that while both Ccl2 and Cxcl10 elicit morphological changes in microglia, they may play differential roles in microglia activation. Notably, both Ccl2 and Cxcl10 are upregulated in aging and AD pathogenesis, suggesting their signaling activity via DSB-bearing neurons could have significant effects on neuroinflammation^44,58,59^. Together, our data indicate Ccl2 and Cxcl10 play important roles in recruiting and activating microglia to neurons burdened with DSBs. This establishes a novel role for neuronal communication with microglia in the context of age-associated neurodegenerative disease, and uncovers a new facet of DSB toxicity in mature postmitotic neurons.

## Discussion

Substantial evidence supports a causal role for microglia and neuroinflammation in AD pathogenesis^29,60,61^. For example, many AD risk genes and genomic loci are most active in microglia^62–64^, and reactive microglia are thought to create a cytotoxic environment for neurons and exacerbate neurodegneration^65–68^. Here, we provide evidence that neurons participate in this inflammatory signaling as well. Specifically, upon the accumulation of DSBs, neurons are able to engage microglia through the secretion of proinflammatory factors, thus facilitating neuroinflammation and disease progression. This is a previously undefined role for neurons in the context of neurodegenerative disease, and provides a mechanistic link between genomic fragility and senescence in neurons and microglia activation.

Interestingly, the activation of inflammatory signaling in DSB-bearing neurons corresponds with a progressive erosion of cell identity. Using single-nuclei RNA-seq, we find that DSB-bearing neurons engage DNA repair pathways first, but transition to immune gene expression at later stages of genome toxicity. These cells are most enriched for senescent markers, such as *Cdkn1a* and *Ubb*, rather than neuronal markers. The age-dependent erosion of neuronal identity has been observed in both sporadic and familial Alzheimer’s disease patients, and coincides with the activation of a number of biological pathways identified in our study, including DNA damage response, cell-cycle re-entry, and NF-kB signaling^31,32^. Interpreted in the context of the present study, these observations could suggest that the age-associated accumulation of DSBs degrades chromatin integrity, ultimately resulting in the activation of immune signaling that engage microglia.

The SASP and canonical antiviral response is activated through the detection of non-endogenous nucleic acids within the cytosol. Thus, it is likely that loss of nuclear and genomic integrity may lead to the release of DNA fragments in DSB-bearing neurons. In fact, we observed increased expression of nucleic acid sensors in DSB-bearing neurons, including cGAS, which plays a significant role in innate immune activation associated with senescence^18^. Nevertheless, many other mechanisms may be at play, including de-repression of transposable elements^69^, epigenetic drift^14^, and ATM-mediated NFkB transcription^19,20^. There is also evidence that nucleic acids released from mitochondria play roles in neuron immune gene expression^70^. It remains to be addressed whether DSB-induced immune gene expression is a product of age-associated decline in DNA repair, or whether multiple cellular functions that are also known to decline with age engage immune signaling as a convergent mechanism of neuronal distress.

We identified NFkB as a major regulator of immune gene expression in DSB-bearing neurons from both CK-p25 mice, primary neurons, and postmortem human brain. In addition, we found that suppression of NFkB activity through p65 knock down or small molecule inhibition primarily in neurons was sufficient to reduce activated microglia morphology and gene expression. NFkB has been previously identified as a therapeutic target for AD, albeit in the context of microglia and astrocytes^71^. Our results now indicate that this transcription factor plays a pivotal role in DSB-bearing neurons as well. Importantly, in rodent models of learning and memory, NFkB activity in neurons is neuroprotective. Suppression of NFkB in forebrain excitatory neurons impairs spatial learning and neuronal plasticity^72^, and exacerbates cell death following exposure to neurotoxic stimuli^73^. Therefore, the primary role of NFkB may regulate healthy neuronal function in times of homeostasis, but it can also orchestrate immune activation in response to cell stressors.

Finally, we identify Ccl2 and Cxcl10 as primary signaling molecules secreted from DSB-bearing neurons to recruit and activate microglia. In the CK-p25 model, DSB-bearing neurons are the first cell type to express Cxcl10 and Ccl2. Other cell types express these chemokines at a later timepoint, presumably in response to DSB-bearing neurons. Furthermore, spatial transcriptomics reveal that signatures of reactive microglia are closely associated with γH2AX-positive neurons, suggesting that DSB-bearing neurons are hubs for neuro-immune communication. Notably, increased levels of both of these cytokines are implicated in the pathogenesis of AD, and affect blood brain barrier permeability to aid the infiltration of peripheral monocytes^23,43,74,75^. However, manipulation of either signaling axis seem to have varying effects on AD pathology. For example, Ccr2 deficiency in murine models of AD aggravate amyloid pathology and cognitive decline, but Ccl2 overexpression seems to also increase amyloid deposition^42,76,77^. Another study demonstrates that deficiency of the Cxcl10 receptor Cxcr3 reduces amyloid deposition and behavioral deficits^78^. These findings suggest that Ccl2 and Cxcl10 are crucial for effective microglia recruitment and activation, but imbalance in these signaling axes have detrimental effects on cognition and pathology clearance.

As a whole, we leveraged bulk, single-nuclei, and spatial transcriptomic techniques in parallel with *in vitro* and *in vivo* manipulations of NFkB signaling to characterize DSB-bearing neurons and their relationship with microglia in the context of age-associated neurodegenerative disease. We demonstrate that DSB accumulation elicits senescent and antiviral-like signaling in neurons, which recruits and activates microglia in an NFkB-dependent manner. Our data posit that neurons play meaningful roles in neuroinflammation, which historically was largely thought to be driven solely by glial cells. Crucially, this axis of neuron-microglia communication is mediated by DNA damage accumulation in neurons, revealing that two hallmarks of Alzheimer’s disease, genome fragility and neuroinflammation, are mechanistically linked.

## Methods

### Fluorescence-activated nuclei sorting

Frozen cortices were disrupted with a handheld homogenizer in ice-cold PBS with protease inhibitors (cat no. 11836170001, Roche, Basel Switzerland) and RNAse inhibitors (cat no. EO0382, Thermo Fisher Scientific, Waltham MA). Samples were fixed with 1% paraformaldehyde for 10 minutes at room temperature, then quenched with 2.5M glycine for 5 minutes. Nuclei were isolated through dounce-homogenization followed by filtration with a 70uM cell strainer (cat no. 21008-952, VWR, Radnor PA). The following antibodies were used to immunotag nuclei: anti-H2A.X-Phosphorylated (Ser139) antibody conjugated to APC (cat no. 613416, BioLegend, San Diego CA), anti-ATM-phospho (Ser1981), antibody conjugated to PE (cat no. 651203, BioLegend, San Diego CA), anti-NeuN antibody conjugated to Alexa Fluor 488 (cat no. MAB377X, EMD Millipore, Burlington MA), anti-PU.1 antibody conjugated to Alexa 647 (cat no. 2240S, Cell Signaling Technology, Danvers MA), anti-RFP antibody (cat no. 600-401-379, Rockland Antibodies and Assays, Gilbertsville, PA), and anti-CaMKII-α (6G9) antibody (cat no. 50049S, Cell signaling Technology, Danvers MA). Antibodies were incubated with nuclei in 1% BSA/PBS at 4°C for one hour or overnight. For non-conjugated antibodies, samples were washed once with 1% BSA/PBS, then resuspended with 1:1000 Alexa Fluor secondary antibody (Thermo Fisher Scientific, Waltham MA) for one hour at 4°C. Samples were strained through a 40um filter (21008-949, VWR) and stained with DAPI (cat no. D9542, Sigma Aldrich, St. Louis MO) before sorting. Sorting was performed on a FACSAria at the Koch Institute Flow Cytometry Core (BD Biosciences, US). At least 50,000 nuclei of each cell type was collected for RNA-sequencing. Nuclei were sorted into 1% BSA/PBS, then spun at 2kG for 15 minutes in a cooled centrifuge (cat no. 97058-916, Epindorf North America) for downstream analysis. For single-nucleus RNA-sequencing, nuclei were not fixed. Single nuclei were sorted into a 96-well plate then transported immediately to the MIT BioMicro Center for library preparation. Flow cytometry analysis was performed using FlowJo software (Ashland, OR).

### Human γH2AX fluorescence-activated nuclei sorting

Frozen temporal cortex tissue samples were homogenized and stained following the protocol described above. A two-week induced CK-p25 mouse sample was used to set a γH2AX-positive/negative threshold for human nuclei.

### Bulk RNA sequencing

For bulk RNA-sequencing of sorted nuclei, samples were treated for 15 minutes with Proteinase K at 50°C and then for 13 minutes at 80°C. RNA was extracted using Direct-zol RNA MicroPrep kit according to manufacturer’s instructions (cat no. R2062, Zymo Research, Irvine CA). Purified RNA samples underwent fragment analysis at the MIT BioMicro Center. Libraries were generated from samples passing quality control (DV200 < 50%). Library generation was performed using the SMARTer Stranded Total RNA-Seq Kit v2 - Pico Input Mammalian (cat no. 634412, Takara Bio Inc), then submitted to the MIT BioMicro Center for quality control (fragment analysis and qPCR), followed by sequencing. Paired-end sequencing was performed using the Illumina NextSeq500 platform according to standard operating procedures.

For bulk RNA sequencing of etoposide-treated primary neurons, cultures were collected in Trizol LS (cat no. 10296028, Thermo Fisher Scientific, Waltham MA). mRNA was purified and extracted using Direct-zol RNA MicroPrep kit according to manufacturer’s instructions. RNA was submitted to the MIT BioMicro Center for library preparation and sequencing. Libraries were prepared using NEBNext^®^ Ultra^™^ II RNA Library Prep Kit for Illumina^®^ (cat no. E7770, New England Biolabs, Ipswich MA). Single-end sequencing was performed using the Illumina NextSeq500 platform according to standard operating procedures.

### Mouse Bulk RNA-seq Read cleaning and bulk RNA-Seq pipeline

From the paired-end fastq files, the first three nucleotides were trimmed off the second sequencing read using cutadapt version 1.16. TrimGalore version 0.4.5 was used in paired mode to trim adapters and low-quality portions of reads. Reads were aligned using Salmon version 0.12.0 to mouse genome version GRCm38.94 and human genome version GRCh38.94. For downstream analysis, TPM files from Salmon were imported into R version 3.6.1 using tximport version 1.14.0 with the option to generate estimated counts using abundance estimates scaled up to library size, and additionally scaled using the average transcript length over samples and the library size (countsFromAbundance = lengthScaledTPM).

### Mouse Bulk RNA-seq Differential Analysis

We performed differential analysis using R version 3.6.1 and DESeq2 version 1.26.0. For each corresponding cell type and condition, we performed differential expression using only the corresponding samples. We selected genes as significant if they met the cutoff threshold (absolute value of the log2 fold change > 1, adjusted p-value < 0.05).

### Mouse Bulk RNA-seq Gene Set Enrichment Analysis

We used the complete set of RNA-Seq results for each differential analysis for downstream GSEA processing. Genes for which adjusted p-value could not be calculated were excluded. We ran GSEA version 3.0 in ranked list mode with default settings. The gene ontology biological pathways gene sets from Molecular Signatures Database v7.3 (MSigDB) were used for analysis. Genes were ranked by the sign of the fold change times the negative base 10 log of the adjusted p-value.

### Enrichr

Significantly upregulated genes from the Stage 2 vs. Baseline contrast were filtered based on their occupation in MSigDB gene ontology biological processes containing the keyword “immune.” This resulted in a Stage 2 immune module comprised of 940 genes. The corresponding Entrez gene symbols were then entered in to the Enrichr website: https://maayanlab.cloud/Enrichr/#. The resulting transcription factor enrichment data were downloaded as tables from the following transcription factor libraries: ChEA 2016, ENCODE and ChEA Consensus TFs from ChIP-X, TRRUST Transcription Factors 2019, Enrichr Submission TF-Gene Coocurrence, TRANSFAC and JASPAR PWMs, and ENCODE TF ChIP-seq 2015. Data are available in Supplementary Table 1. More information about the Enrichr libraries can be accessed on the Enrichr website.

### Smart-seq pipeline

#### SMART-seq2

Single nuclei library preparation was performed by the MIT BioMicro Center using the SMART-Seq® v4 Ultra® Low Input RNA Kit for Sequencing (Takara Bio Inc) according to manufacturer’s instruction. Libraries were sequenced on a MiSeq Illumina sequencer according to standard operating procedures.

#### Read processing

We aligned 40-bp paired-end reads to the mm10 genome for each of 1,357 single nuclei from 12 mice (64 cells each for 6 CK controls and an average of 162 cells in each of 6 CKp25 mice) using bwa mem (options: -k 15 -M) and filtered out improperly aligned reads and secondary alignments with samtools (options: -f 3 -F 1280). We ran HTSeq-count10 on each cell’s filtered bam file to compute the cell’s transcriptomic coverage over each gene’s exons in vM25 GENCODE gene annotation.

#### Transgene detection

We first aligned 40-bp single-end reads from each sequenced single nucleus separately with the STAR aligner6 against the b37 genome with decoy contigs using a two pass alignment (options: - -outFilterMultimapNmax 20 --alignSJoverhangMin 8 --alignSJDBoverhangMin 1 -- alignIntronMin 20 --alignIntronMax 1000000). We then used Picard tools7 to revert and merge the alignment with unaligned reads and marked duplicates on the merged bam file. We identified and removed alignments on decoy contigs, sorted and fixed NM, MD, and UQ tags with Picard tools, filtered duplicates, unmapped, and non-primary alignment reads, and split reads by Ns in their CIGAR string.

### Cell type annotation

#### Cell identities

We used SCANPY11 to process and cluster the expression profiles and infer cell identities. We kept only 21,859 protein coding genes detected in at least 3 cells and filtered out 349 cells with less than 100 expressed genes, leaving 1,008 cells over the 12 mice. We used the filtered dataset to calculate the low dimensional embedding of the cells (UMAP) from the log1p matrix PCA with k=50 and nearest neighbor graph with N=20 (UMAP default parameters, min_dist=0.2), and clustered it with Leiden clustering (resolution=2), giving 17 preliminary clusters. We then manually assigned clusters based on the following 2-3 major marker genes per class: Excitatory neurons: Camk2a, Gria2, Syt1; Inhibitory neurons: Gad1, Gad2; Astrocytes: Gfap; Microglia: Cd33, Csf1r; Oligodendrocytes: Plp1, Mbp; Oligodendrocyte progenitor cells (OPCs): Bcan; Stage 2: Cdkn1a, Ubb. We merged clusters sharing marker genes to obtain 7 final clusters, defining two broad neuronal subtypes (484 excitatory and 108 inhibitory cells), three glial clusters (50 microglia, 131 oligodendrocytes, and 33 OPCs), 179 Stage 2 cells, and 23 cells with both high read counts and broad, non-specific marker gene expression, likely due to doublets or other sorting artifacts. The doublet cluster was removed from downstream analyses. We also identified a cluster of cells that expressed fewer than 500 genes and had high expression of mitochondrial genes. Due to the questionable quality of these cells, this cluster was also removed from downstream analysis. We further sub-annotated neuronal subtypes from neuronal clusters with distinctive expression in the original Leiden clustering, resulting in four excitatory subclusters (314 Ex0, 91 Ex1, 32 Ex2, and 47 Ex3 neurons) and two inhibitory subclusters (71 In0 and 37 In1 neurons).

#### Signature analysis on single-cells

For each signature, a joint expression value was calculated as the number of reads per 10k reads in each cell coming from all of the signature’s genes (genes with adjusted p-values < 0.05). Signature expression was transformed by log1p and averaged across all cells in a given neuronal subtype and mouse to obtain average signature values for plotting. We used two-sided t-tests to compare the signature expression levels of pairs of excitatory neuronal subtypes (Supplementary Figure 3f-h).

### Trajectory analysis

#### Pseudotime

We performed pseudotime analysis using Monocle3 (v0.2.3.0) on the 349 Ex1, Ex2, Ex3, and Stage 2 neuronal populations. We normalized total counts per cell in the read count matrix to the median number of counts, log1p transformed the matrix, and regressed out the counts per gene using Monocle. We clustered the subsetted data in a new UMAP, clustered cells, learned a trajectory graph, and ordered cells by choosing the initial node as the Ex1 end of the graph to get a pseudotime across the graph. We plot the log1p read counts across the gene set as well as the generalized additive model (GAM) smoothed fits of each signature’s per-cell expression values across pseudotime (Figure 2e,f).

#### Differential genes on pseudotime

We used tradeSeq to detect differentially expressed genes along the pseudotime, by fitting GAMs to each gene along pseudotime with uniform weights on each cell and performing an association test to test for genes with pseudotime-associated expression patterns. We found 530 significant genes. We selected a number of representative genes and plotted their smoothed fit (binned into 100 bins across pseudotime and scaled) and their expression (log1p per-cell normalized counts) across the 349 pseudotime-ordered cells. We labeled each gene with the neuronal subtype closest to the pseudotime point with its peak expression (Supplementary Figure 3f).

#### Ontology enrichment

We created 20 tiled windows of 20 quantiles in width across cellular pseudotime, for each window selected genes whose peak smoothed expression fell into the pseudotime range, and performed GO enrichment on each window’s genes using gprofiler, limiting results to terms with fewer than 2,500 genes in GO and Reactome. For visualization, we manually curated 34 representative top enriched terms across all Reactome and GO enrichments (Supplementary Figure 3j).

#### Human bulk RNA-seq analysis

Paired-end reads were trimmed for adapters using TrimGalore (https://github.com/FelixKrueger/TrimGalore) and aligned to the human genome reference GRCh38/hg38 using the R software package Rsubread (Liao, Smyth, and Shi 2019). Mapped reads were summarized to gene level counts using the featureCounts function of Rsubread, considering the latest RefSeq Reference Genome Annotation for gene reference. Protein coding genes with detected counts in at least 1 sample library were retained and normalized in rpkm units. Normalization and differential expression analysis was performed using the edgeR package (Robinson, McCarthy, and Smyth 2010). All statistical analyses and plots were performed using the statistical programming language R (http://www.r-project.org/). Gene ontology and functional enrichment analyses were performed using gprofiler2 package (Kolberg et al. 2020).

#### Human bulk gene set activity analysis

Gene set activity for cell type marker genes and gene signatures identified in the Cpk25 mouse model were quantified per human bulk RNAseq sample using Gene Set Variation Analysis (GSVA) (Hänzelmann, Castelo, and Guinney 2013), as implemented in the Bioconductor package GSVA (https://www.bioconductor.org/packages/release/bioc/html/GSVA.html). For cell cell type signature analysis the consensus human brain cell type markers reported in (Mohammadi, Davila-Velderrain, and Kellis 2020) were used as reference. Differential gene signature activity across conditions (γH2AX +/ctr, AD/noAD) and/or cell groups (NeuN+/NeuN-) was assessed using a two-sided wilcoxon rank sum test.

#### Human snRNA-seq analysis

Single-nucleus transcriptomic sequencing data from postmortem cortical samples (prefrontal cortex, Brodmann area 10) of 48 subjects with varying levels of AD pathology was obtained from (Mathys et al. 2019). Individual-level celltype expression profiles were computed by averaging for each individual the normalized gene-expression profiles across cells of the same cell type. Average profiles were subsequently mean-centered and scaled to compute gene-wise correlation coefficients of gene expression versus individual-level measures of global AD pathology burden reported as part of the ROSMAP cohort. Briefly, global burden of AD pathology is a quantitative summary of AD pathology derived from counts of three AD pathologies: neuritic plaques (n), diffuse plaques (d), and neurofibrillary tangles (nft), as determined by microscopic examination of silver-stained slides (Mathys et al. 2019).

Global consistency between gene signatures observed in CKp25 mice and a neuronal-specific association between gene expression and AD pathology in human tissue was assessed statistically using a nonparametric resampling test. To test whether cell-type specific expression of CKp25 Stage 2 signature genes tends to correlate with pathology in the human brain, a z-score statistic was computed to quantify the deviation of their correlation coefficient rank scores, relative to random expectation. Expected scores were estimated by randomly sampling same-sized gene sets (n = 1,000 replicates). This analysis was performed for excitatory neurons, inhibitory neurons, and microglia cells independently.

#### Stage 1 and 2 signature generation

Stage 1 and Stage 2 gene signatures were curated by performing differential expression analysis on the CK-p25 bulk RNAseq dataset. Stage 1 vs Baseline provided the genes for the Stage 1 signature, and Stage 2 vs Baseline provided the genes for the Stage 2 signature. Only genes which met the cutoff threshold (log2 fold change > 1, adjusted p-value < 0.05) were retained.

#### Visium spatial transcriptomics library generation

Mice were transcardially perfused with ice-cold saline, then brains were dissected and flash frozen in OCT. A cryostat was used to generate 10uM coronal sections of the hippocampus. These sections were applied to 10X Visium Spatial Gene Expression slides. Sections were immunostained with γH2AX and DAPI following manufacturer’s instructions. Sectioning and staining was performed at the MIT Hope Babette Tang (1983) Histology Core Facility. Sections were imaged immediately after staining using a Olympus FV1200 Laser Scanning Confocal Microscope at the MIT Microscopy Core Facility. Sections were then used to generate 10X Visium Spatial Gene Expression Libraries according to manufacturer’s instructions at the MIT BioMicro Center. Libraries were sequenced on a NovaSeq6000 Illumina sequencer according to standard operating procedures.

#### Visium spatial transcriptomics data processing

Samples were processed using Scanpy 1.7.2. The 7 sample data matrices were merged into one matrix which was then processed. The sample id and location of each capture area of the resulting matrix were saved and used for visualization. Counts were normalized (total count of 10,000 per capture area) and logarithmized (using scanpy’s log1p function). The resulting counts matrix, called raw normalized was used for expression visualization and differential expression analysis. For dimension resolution purposes, the raw normalized matrix was further processed: genes that were not characterized as highly variable enough were filtered out (minimum mean of 0.0125, max mean of 3, minimum dispersion of 0.5), and linear regression was performed to eliminate the effect of covariates (total_counts, and mitochondrial genes percentage). The data was then scaled (standard scaling, max value of 10). Afterwards, PCA was performed, as well as sample-level batch correction, using Harmony. Then, a knn network was constructed for the creation of a UMAP embedding. Clusters were discovered using the leiden algorithm.

#### Visium immunohistochemistry image processing

Immunostaining images were first processed as shades of gray pictures. The largest autofluorescence artifacts were removed manually. The signal was then amplified and cleaned using a 85% contrast increase on each image.

As the immunochemistry images of the tissue align perfectly to the pictures taken for 10X Visium purposes, calibration was performed to reconstruct the capture areas grid on the immunostaining images. In that image, for each capture area, the mean signal within a circle maximizing image coverage is calculated, and recorded as the DNA-Damage signal. After standard scaling of this variable, a threshold of 0.4 standard deviation was set, to assign a capture areas as positive or negative for DNA damage.

#### Visium differential expression and Mathys et al., 2017 microglia signature analysis

Differential expression analysis was performed using Wilcoxon rank sum test, and the resulting p-values were corrected using Benjamini-Hochberg FDR correction. For GSEA, the package gseapy was used, with 100 permutation and the signal-to-noise method. As input, the raw normalized matrix was used, though only containing the genes considered as highly variable in the dataset.

#### RT-qPCR

RNA was extracted from primary tissue cultures using the RNeasy Plus mini kit (cat no. 74136, Qiagen, Hilden Germany). Reverse transcription was performed using Invitrogen SuperScript IV First Strand Synthesis System with Oligo dT primers according to the manufacturer’s protocol (cat no. 18091050, Thermo Fisher Scientific, Waltham MA). cDNA was quantified with a NanoDrop spectrophotometer. qPCR was performed using a Bio-Rad CFX-96 quantitative thermal cycler (cat no. 1855195, Bio-Rad, Hercules CA) and SsoFast EvaGreen Supermix (cat no. 1725202, Bio-Rad, Hercules CA). Relative changes in gene expression were determined using the 2^−ΔΔCt^ method. Cycle numbers for the gene *Gapdh* or *Rpl11* were used for housekeeping Ct values.

#### Immunofluorescent microscopy

Mice were transcardially perfused with ice-cold PBS, then fixed with ice-cold 4% paraformaldehyde in PBS. Dissected brains were drop-fixed overnight in 4% paraformaldehyde in PBS at 4°C. Forebrains were sectioned with a vibrating microtome (Leica BioSystems, Wetzlar Germany) to generate 40 uM coronal slices. Slices were blocked for two hours at room temperature in blocking buffer (10% NGS, 0.3% Triton X-100, PBS), then incubated with primary antibody overnight at 4°C. Slices were washed 3 x 10 minutes with PBS, and Alexa Fluor Secondary antibodies (Thermo Fisher Scientific) were added at a 1:1000 dilution for 1 hour at room temperature. Slices were washed again 3 x 10 minutes with PBS, then stained with DAPI (cat no. D9542, Sigma Aldrich, St. Louis MO) and mounted onto Fisherbrand™ Superfrost™ Plus Microscope Slides (cat no. 12-550-15, Thermo Fisher Scientific, Waltham MA) with Fluoromount-G™ Slide Mounting Medium (cat no. 100502-406, VWR, Radnor PA).

Primary neurons cultured on cover glass (cat no. 194310012A, VWR, Radnor PA) were washed once with PBS, then fixed with 4% paraformaldehyde/PBS for 15 minutes at room temperature. Immunostaining proceeded as described.

Free-floating slices, at 40um thickness from postmortem human brain blocked for 1 hour at room temperature in blocking buffer before being incubated in primary antibodies for 72 hours at 4°C with gentle rocking in blocking buffer (Anti-NeuN, Synaptic Systems cat no. 266 004, 1:500; Anti-phospho-Histone H2A.X, Millipore cat no. 05-636, 1:100; Anti-NFkB p65 Invitrogen cat no. 51-0500, 1:300). Samples were then rinsed three times in 1xPBS for 5 minutes and incubated in secondary antibodies (Alexa Fluor-488, 594 or 647, ThermoFisher Scientific; 1:1,000) for 2 hours at room temperature. After 1x PBS rinse, samples were incubated in 1:10,000 Hoechst in PBS (Invitrogen cat no. H3569) followed by 2 minutes in TrueBlack Lipofuscin Autofluorescence Quencher (Biotium cat no. 23007) with 3 subsequent 1x PBS rinses before mounting and imaging.

Mounted samples were imaged with a Zeiss LSM 710 confocal microscope. Images were quantified using ImageJ (NIH Image Analysis) and Imaris (Oxford Instruments). At least two coronal slices were used for each mouse for image quantification. For PHP.eb AAV shp65-RFP cell-type analysis, 40 uM coronal brain sections were stained for RFP and either NeuN, Iba1, GFAP, or Olig2. First, RFP-positive cells were identified in each image. Then, the percent of RFP-positive cells that also had positive immunoreactivity for a given cell-type marker was quantified. GFAP-positive cell number and intensity were calculated using Imaris software (Oxford Instruments, UK).

#### Primary Neuron Culture

Cortices were dissected from E15 Swiss-Webster embryos in ice-cold HBSS (cat no. 14175103, Thermo Fisher Scientific, Waltham MA) and dissociated with papain (cat no. LS003126, Worthington Biochemical Corp., Lakewood NJ) and DNAse I (cat no. 10104159001, Roche, Basel Switzerland). Cells were resuspended in plating media (Neurobasal media (cat no. 21103049, Thermo Fisher Scientific, Waltham MA), 1% Penicillin/Streptomycin Solution (cat no. 400-109, Gemini Bio-Products, Sacramento CA), 10% FBS)) and filtered through a 100 uM cell strainer (cat no. 21008-950, VWR, Radnor PA). Cell density was quantified using a Countess II Automated Cell Counter (cat no. AMQAX1000, Thermo Fisher Scientific, Waltham MA), then plated on poly-D-Lysine-coated 12-well culture dishes, 0.5×10^6. Cultures were maintained in 5% CO2 at 37 °C in a cell culture incubator. After allowing four hours for the cells to adhere to the plate, the media was replaced and maintained with neurobasal media supplemented with B-27 (cat no. 17504-044, Invitrogen, Carlsbad CA), 1% Penicillin/Streptomycin, and 1% GlutaMAX Supplement (cat no. 35050-079, Thermo Fisher Scientific, Waltham MA).

#### Etoposide Treatment

Primary cortical neuron cultures (DIV11-13) were treated with 1, 5, 10, 25, or 50uM etoposide prepared from 20mM stock (cat no. E1383-250MG, Sigma, St. Louis MO). Cultures were treated for either 3 or 6 hours before collection for downstream experiments. We used two SASP cytokines, *Ccl2* and *Cxcl10* (found significantly upregulated in Stage 2 neurons) as biomarkers for activation of immune signaling. Quantitative reverse transcription PCR (RT-qPCR) was used to measure gene expression. A six-hour 50 uM etoposide treatment resulted in a robust induction of compared to other time points and drug concentrations. We also observed that a 50uM etoposide treatment resulted in a marked increase in γH2AX nuclear intensity compared to control cultures (Figure 2a).

#### NFkB Activation Inhibitor

Primary neuron cultures were treated with 10uM NF-kappaB Activation Inhibitor VI, benzoxathiole compound from Abcam (cat no. ab145954). Cultures were pre-treated for at least 30 minutes before ETP exposure.

#### RNAScope In-Situ Hybridization

Fluorescent in-situ hybridization was performed using the RNAscope® Multiplex Fluorescent Reagent Kit v2 according to manufacturer’s instructions (cat no. 323100, Advanced Cell Diagnostics, Newark CA). Probes targeting murine *Ccl2* (cat no. 311791) and *Cxcl10* (cat no. 408921) were purchased from Advanced Cell Diagnostics. Following the RNAscope protocol, slices were stained for γH2AX (cat no. 05-636, EMD Millipore, Burlington MA). Samples were imaged with a Zeiss LSM 710 confocal microscope at 40x objective. Images were analyzed with ImageJ.

#### RNAscope Analysis

ImageJ version 2.1.0 thresholding was used to identify γH2AX-positive nuclei and generate ROIs. Process -> “Find Maxima” was used to count mRNA puncta within ROIs. Prominence >100.00 with “Strict” setting. The number of γH2AX-positive nuclei with 2≤ maxima were quantified, as well as the number of maxima per γH2AX-positive nucleus. γH2AX-negative nuclei were identified by thresholding for individual nuclei using the DAPI channel, then excluding ROIs that overlapped with γH2AX-positive ROIs. Individual nuclei were identified following the Nuclei Watershed Separation process described on the ImageJ website: https://imagej.net/Nuclei_Watershed_Separation

### Brain tissue samples

#### MADRC brain tissue samples

Fresh frozen postmortem brain samples were generously provided by the Massachusetts Alzheimer’s Disease Research Center. These samples were used for bulk RNA-sequencing. Individuals were selected based on clinical diagnosis and Braak score. The three samples labeled as AD all had a clinical diagnosis of AD, and a Braak score of VI. The three samples labeled as non-AD did not have a clinical diagnosis of AD, and had Braak scores of I, II and II. Sample metadata is available in Supplementary Table 3.

#### ROSMAP brain tissue samples

Fixed frozen postmortem brain samples were chosen from the Religious Orders Study and Memory and Aging Project cohort (ROSMAP). ROSMAP is a longitudinal cohort study of ageing and dementia in elderly nuns, brothers and priests. Sample metadata is available in Supplementary Table 4. In-depth description of metadata variables are available at the Rush Alzheimer’s Disease Center (RADC) website: https://www.radc.rush.edu/docs/var/variables.htm.

#### P65 knock-down

Custom p65 and scramble shRNA oligos cloned into an AAV backbone (pAV-U6-RFP) were purchased from ViGene Biosciences (Rockville, MD). P65 shRNA sequences were:

1. GATCCGGCAGGCTATCAGTCAGCGCATTGTGCTTATG CGCTGACTGATAGCCTGCTTTTTA
2. GATCCGCGGATTGAGGAGAAACGTAAATGTGCTTTTTA CGTTTCTCCTCAATCCGTTTTTA
3. GATCCGCACCATCAACTATGATGAGTTTGTGCTTAACT CATCATAGTTGATGGTGTTTTTA
4. GATCCGCCTGAGGCTATAACTCGCCTATGTGCTTTAGG CGAGTTATAGCCTCAGGTTTTTA
5. shRNA scramble: GATCCGCAACAAGATGAAGAGCACCAACTCGAGTTGGTGCTCTTCATCTTGTTGTTTTTA

P65 knockdown was confirmed via RT-qPCR in-house. PHP.eB AAV was generated by Janelia Viral Services. PHP.eB AAV shRNA or PBS was delivered retro-orbitally to anesthetized CK-p25 mice, 2×10^11. Two weeks after injection, mice were induced by removing doxycycline diet. Following induction, mice were transcardially perfused with ice-cold PBS. One hemisphere was drop-fixed overnight in 4% paraformaldehyde/PBS at 4°C for immunostaining. The other hemisphere was flash-frozen in liquid nitrogen and stored at −80°C for RNA sequencing. 5 Scramble and 5 p65 knock-down CK-p25 mice were used for RFP-positive γH2AX-positive RT-qPCR. A separate cohort of 7 CK, 5 Scramble, and 6 p65kd CK-p25 mice were used for PU.1-positive RNA-sequencing, and Iba1 and GFAP imaging analyses.

#### CK-p25 mice

CK-p25 double transgenic mice were raised and maintained on a doxycycline diet. All mice were induced by removing doxycycline from their diet to drive the expression of p25-GFP in forebrain excitatory neurons. All mice were induced at 3-4 months old.

#### Conditioned media and Immunodepletion

Following etoposide treatment, cultures were washed once with PBS. Cultures then recovered in fresh media for 24 hours. Media was collected and spun at 2,000g for 10 minutes to remove cellular debris. Media was stored at −80°C for future experiments. Conditioned media from etoposide-treated neurons were incubated with IgG, Ccl2, or Cxcl10 antibodies (all 40ug/mL) for 4 hours at 4°C. Dynabeads™ Protein G were added to pull down the antibody complex.

#### Organotypic Brain Slice Culture

8 through 12 week-old Cx3cr1-GFP male mice were anesthetized with isoflurane and transcardially-perfused, dissected, and sliced in ice-cold NMDG-cutting solution containing (in mM): 2.5 KCl, 0.5 CaCl2, 10 MgSO4, 1.25 NaH2PO4, 20 HEPES, 2 Thiourea, 5 sodium ascorbate, 3 sodium pyruvate, 92 NMDG, 30 NaHCO3, 25 D-Glucose, pH 7.3 – 7.4 with HCl. The slicing chamber was bubbled with 95% O2/5%CO2, and coronal slices were cut at 250 µm thickness using a vibratome (Leica, VT1000s). After the last slice was collected, slices were transferred to a well-plate containing fresh aCSF and placed in an incubator set at 37□C and 95% O2/5% CO2 for 30 minutes. The aCSF solution contained (in mM): 125 NaCl, 2.5 KCl, 1.2 NaH2PO4, 1.2 MgCl2, 2.4 CaCl2, 26 NaHCO3, 11 D-Glucose. Afterward, 100% of aCSF was removed and a 1:1 mixture of fresh aCSF and conditioned media was added to the slices and placed back into the incubator for 6 hours. At the completion of the experiment, slices were fixed overnight at 4°C with 4% paraformaldehyde. Slices were then incubated in 30% glucose overnight at 4°C. Slices were sub-sectioned into 25um slices using a cryostat, then cover-slipped for imaging.

#### ELISA

The Mouse MCP1 ELISA Kit and Mouse IP-10 ELISA Kit from Abcam (cat no. ab208979 and ab214563 respectively) were used to quantify Ccl2 and Cxcl10 in conditioned media from primary neurons. Assays were quantified on a plate reader and protein concentration was calculated according to manual instructions.

#### Microglia morphological analysis

Microglia morphology from CK-p25 and acute slice cultures was analyzed according to the protocol described in Young and Morrison, 2018 with minor alterations^79^. Gray Scale Attribute Filtering (default settings, connectivity: 8) from the MorphoLibJ plug-in version 1.4.1 was used to reduce background noise when thresholding images for skeleton analysis. Following skeleton analysis, Morphological Filters (Operation: Opening, Element: Octagon, Radius (in pixels): 2) from MorphoLibJ was used to quantify soma area. This analysis was performed for both Iba1 and Cx3cr1-GFP imaging experiments.

## Acknowledgements

We thank M. Saturno-Condon, M. Jennings, M. Griffin, and G. Paradis from the Swanson Biotechnology Center Flow Cytometry Facility for assistance with FANS. We thank S. Levine, N. Kamelamela, and A. Hendricks from the MIT BioMicro Center for assistance with RNA sequencing and library preparation. We thank K. Cormier from the MIT Hope Babetter Tang (1983) Histology Core Facility and J. Khun from the MIT Microscopy Core Facility for assistance with Visium Spatial Gene Expression sample preparation. This work was supported by NIH grants AG054012, AG058002, AG062377, NS110453, NS115064, AG067151, AG062335, MH109978, MH119509, and HG008155 (to M.K. and L.H.T), 5 R01 NS102730-03 (to L.H.T), and CureAlz CIRCUITS, and the Glenn Foundation for Medical Research. This work was also supported by NIH fellowship 5F31NS113464-02 to G.W. V.D. is supported by the AARF-19-618751 grant from the Alzheimer’s Association. M.B.V. is supported by HHMI Hanna H. Gray Fellowship. C.A.B. is supported by the NIH training grant GM087237.

---

**Supplementary Figure 1.**
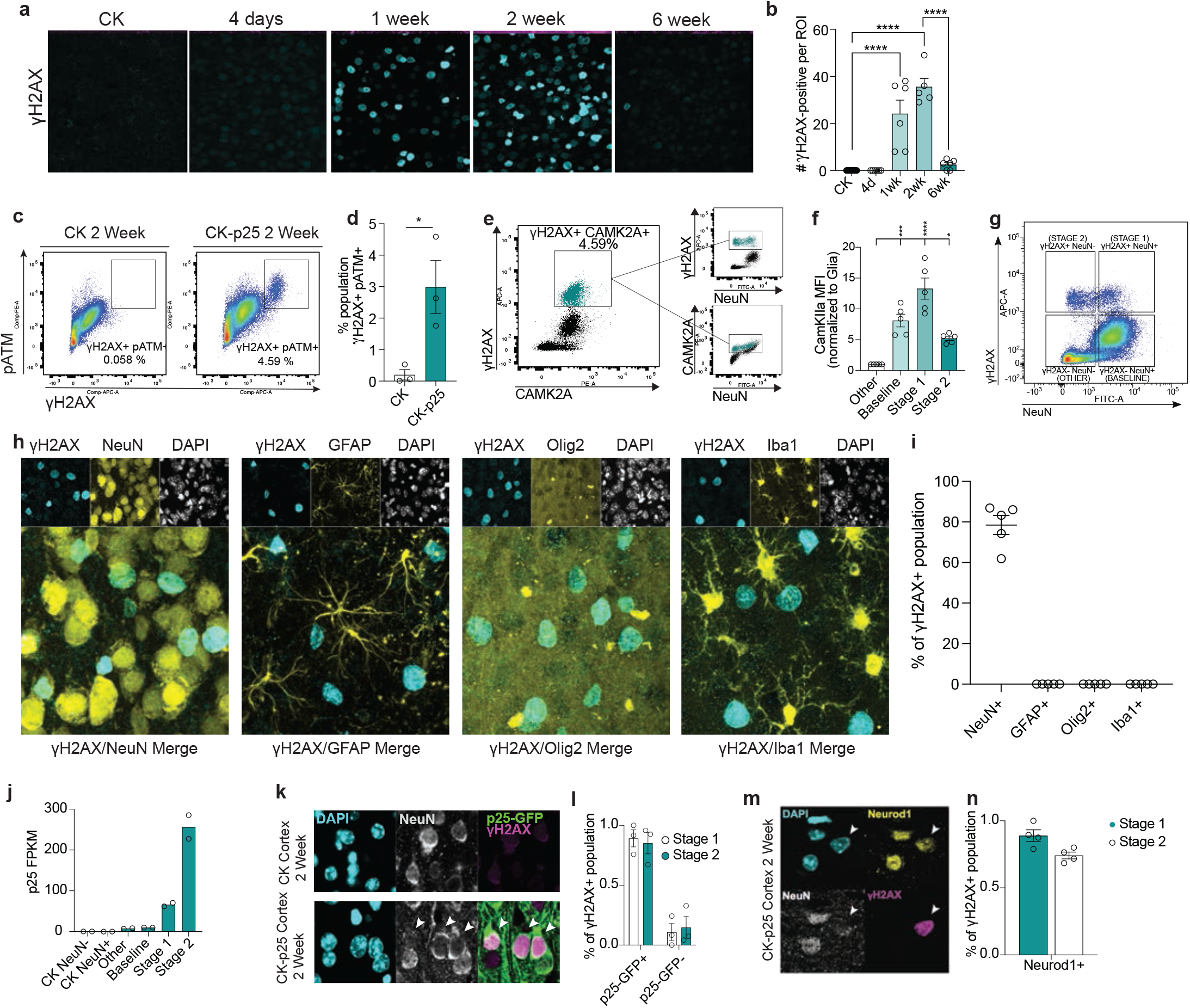
**a.** Representative images of γH2AX immunostaining in CK and CK-p25 cortex over a 6-week timeline analysis. **b.** Average number of γH2AX+ nuclei quantified per image at each timepoint. Each data point represents one mouse. **c.** Flow cytometry dot plot of γH2AX+ pATM+ nuclei in CK and CK-p25 cortex. **d.** Quantificaton of γH2AX+ pATM+ nuclei in CK and CK-p25 cortex. Each data point represents percent γH2AX+ pATM+ nuclei for one mouse. **e.** Flow cytometry dot plot of γH2AX, Camk2a, and NeuN immunoreactivity in nuclei from CK-p25 cortex. γH2AX+ Camk2a+ nuclei are gated in the left graph. This gated population is highlighted in turquoise in the two graphs to the right. **f.** Camk2a median fluorescent intensity (MFI) is quanitifed for Glia, Baseline, Stage 1, and Stage 2 populations. MFI values are normalized to the Glia population. **g.** Representative dot plot of γH2AX and NeuN immunoreactivity in 2-week induced CK-p25 cortex. **h.** Representative images of cell type and γH2AX immunostaining in the 2wk-induced CK-p25 cortex. Celltype markers from left to right: neurons (NeuN), astrocytes (GFAP), oligodendrocytes and oligodendrocyte precursor cells (Olig2), microglia (Iba1). **i.** Quantification of percent of γH2AX-positive nuclei overlapping with each celltype stain. Each dot represents one mouse. 50-100 γH2AX-positive nuclei were analyzed for each mouse and celltype marker. **j.** Expression p25 in CK and CK-p25 gated populations. One datapoint represents fragments per kilobase per million (FPKM) from a gated population from one mouse. **k.** Representative images of GFP, γH2AX, and NeuN immunostaining from CK and CK-p25 cortex at the 2wk time point. Arrowheads indicate γH2AX+ nuclei that express p25-GFP regardless of NeuN immunoreactivity. **l.** Quantification of GFP expression across Stage 1 and Stage 2 populations. Each data point represents percent γH2AX+ population from Stage 1 or Stage 2 from one CK-p25 mouse. **m.** Representative images of Neurod1, γH2AX, and NeuN from CK-p25 cortex at two-week timepoint. White arrowhead indicates a γH2AX+ nucleus with Neurod1 immunoreactivity but not NeuN immunoreactivity. **n.** Quantification of Neurod1 expression across Stage 1 and Stage 2 populations. Each data point represents percent γH2AX+ population from Stage 1 or Stage 2 from one CK-p25 mouse. Error bars represent standard error of mean (S.E.M.); ****P<0.0001, ***P<0.001, *P<0.05, n.s. not significant; One-way ANOVA with Tukey’s test for multiple comparisons (b,f). Student’s T-test (d). Data are averages of 4 images per mouse (b). Data are representative of at least 2 independent experiments (d). Data are pooled from 2 independent experiments (f).

**Supplementary Figure 2.**
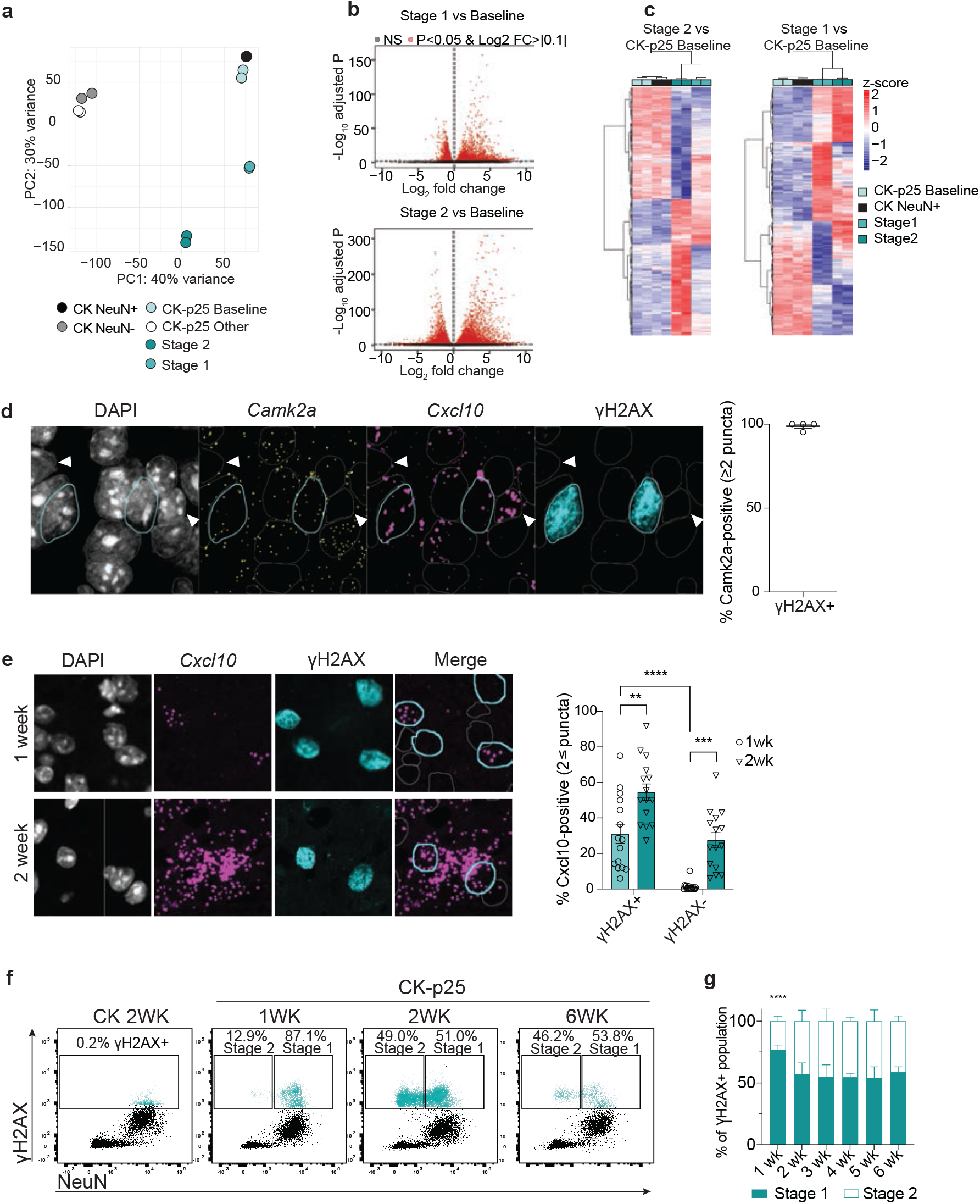
**a.** Principle component analysis (PCA) of normalized gene expression matrix from CK and CK- p25 subpopulations. Principle component 1 (PC1): 66% variance. Principle component 2 (PC2): 29% variance. **b.** Volcano plots of (top) Stage 1 vs. CK-p25 Baseline and (bottom) Stage 2 vs. CK-p25 Baseline contrasts. Gray circles: non-significant (ns) transcripts. Red circles: transcripts with FDR adjusted p-value <0.05 and log2fold change >|0.1|. **c.** Heatmap of differentially expressed genes from Stage 2 vs. CK-p25 Baseline and Stage 1 vs. CK-p25 Baseline contrasts. Each column represents one mouse. **d.** Representative image of RNAscope probes for Camk2a (yellow) and Cxcl10 (magenta), and γH2AX immunostaining (turquoise). To the right, the percent of γH2AX-positive cells that are Camk2a-positive (2 or more Camk2a puncta within the nucleus) are quantified. Each datapoint represents one 2-week induced CK-p25 mouse. 17-42 γH2AX-positive cells were quantified for each mouse. Turquoise outlines indicate γH2AX-positive Camk2a-positive cells. Gray outlines indicate γH2AX-negative cells. White arrows indicate Camk2a-negative cells. **e.** Representative image of the RNAscope probe for Cxcl10 (magenta) combined with γH2AX immunostaining (turquoise). Imaging was performed on 1wk and 2wk CK-p25 cortices. To the right, the number of γH2AX-positive and γH2AX-negative cells with 2 or more Cxcl10 puncta are quantified. Each datapoint represents the average %Cxcl10-positive cells in one image from one mouse. 4-3 images were taken per mouse. (CK-p25 1wk n=4, CK-p25 2wk n=4). **f.** Representative dot plots of γH2AX and NeuN immunoreactivity in CK and CK-p25 mice at 1, 2, and 6 weeks induction. Stage 1 and Stage 2 percent population was calculated with respect to the total γH2AX-positive population. **g.** Stage 1 and Stage 2 percent population are quantified for each timepoint. Error bars represent standard error of mean (S.E.M.); ****P<0.0001, ***P<0.001, **P<0.01, *P<0.05, n.s. not significant; Two-way ANOVA followed by Sidak’s test for multiple comparisons (e,g). Data are representative of 2 independent experiments (d). Data are pooled from four independent experiments (g).

**Supplementary Figure 3.**
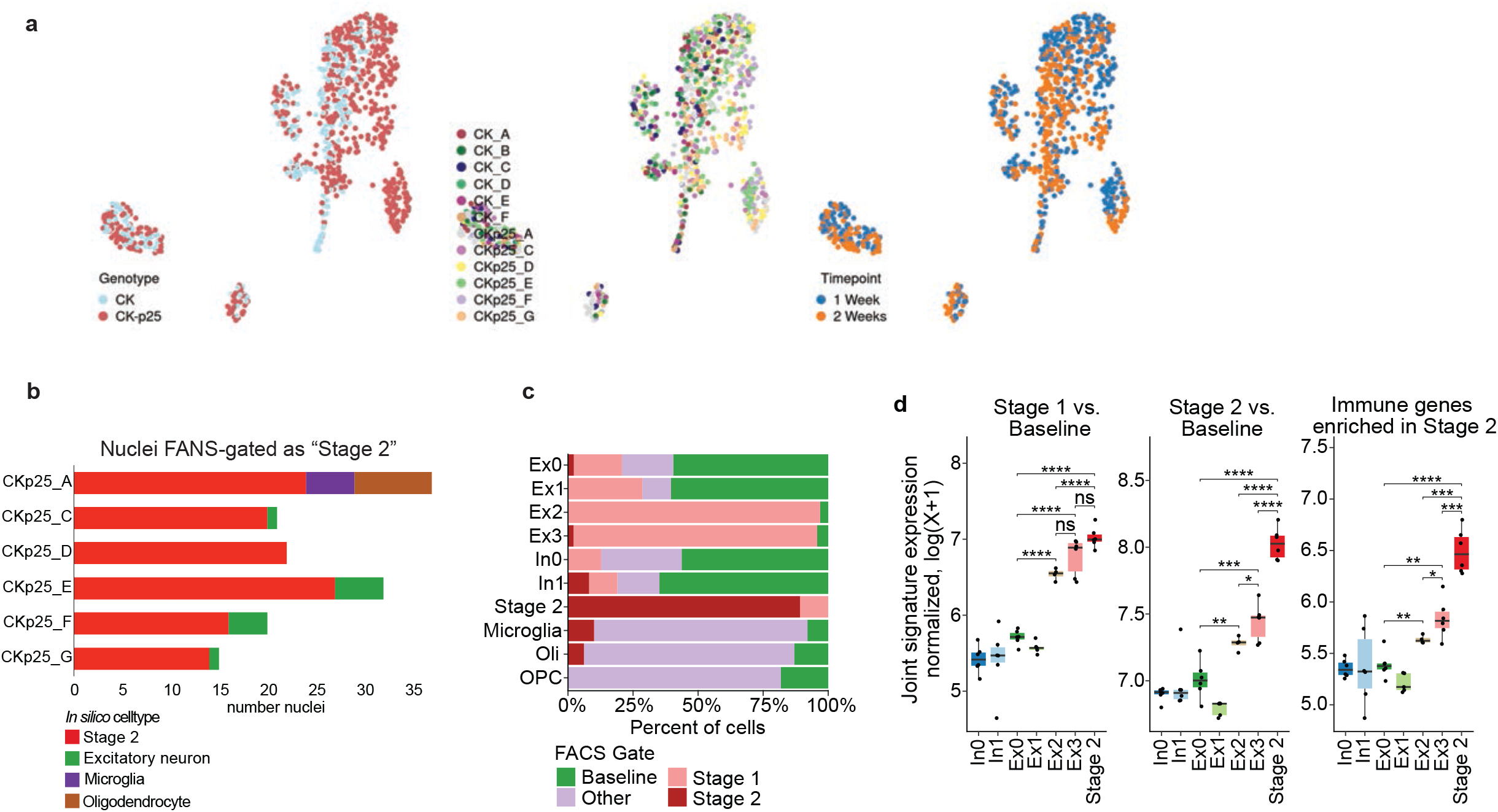
**a.** UMAPs labeled by genotype, mouse id, and timepoint. **b.** Distribution of Stage 2-gated nuclei in in silico cell type clusters, stratified by biological replicate. **c.** Percent FANS label distribution across scRNA cell type clusters. Bargraph colors refer to FANS gate label. **d.** Bulk RNA-seq gene signature enrichment in neuron cell type clusters. Stage 1 signature enrichment (left), Stage 2 signature enrichment (middle), and immune genes from Stage 2 enrichment (right). Each datapoint indicates average gene expression across all cells from one mouse. Only CK-p25 mice were used for this analysis. **e.** ****P<0.0001, ***P<0.001, **P<0.01, *P<0.05, n.s. not significant; One-way ANOVA with Tukey’s test for multiple comparisons (d).

**Supplementary Figure 4.**
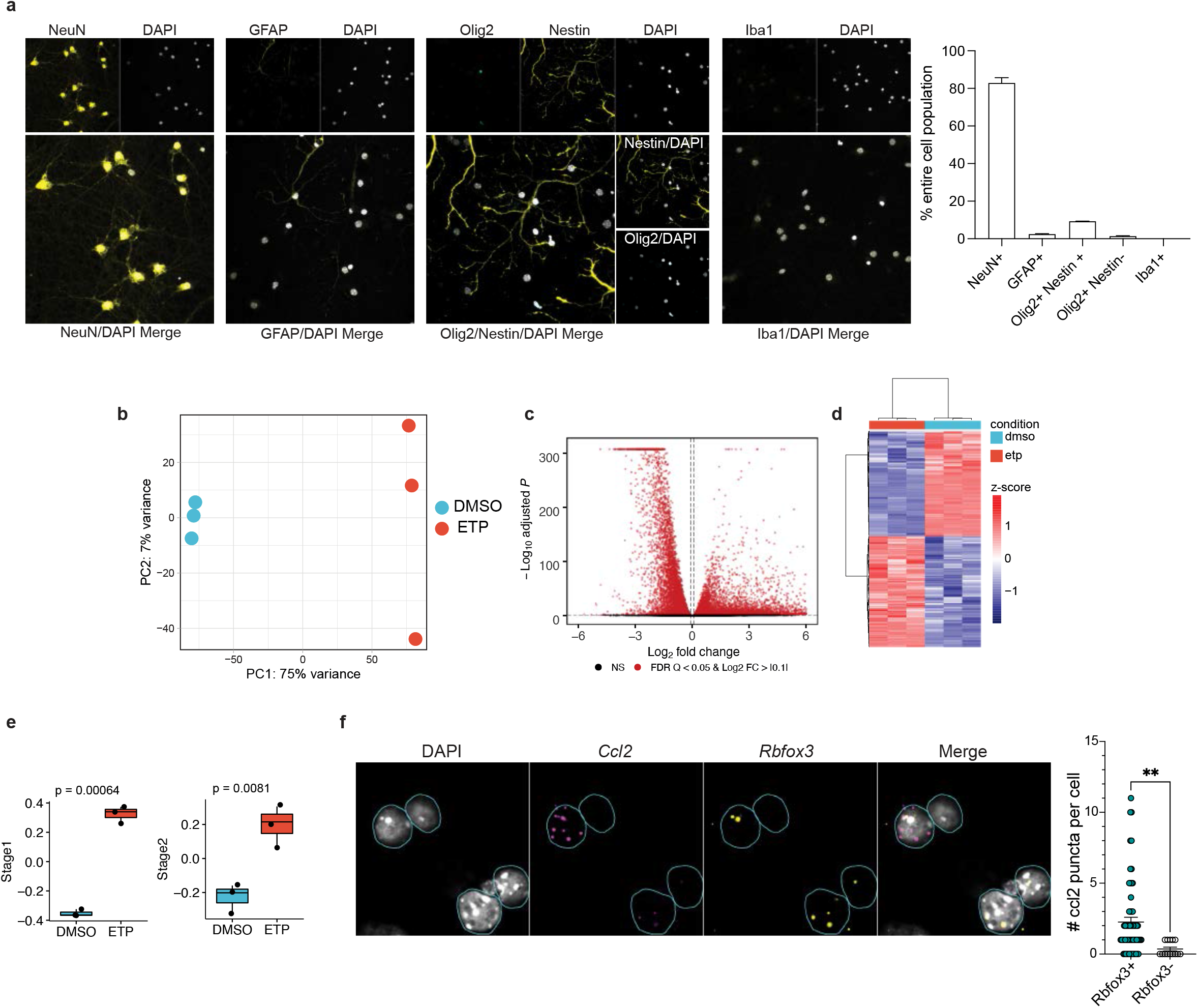
**a.** Representative images of celltype immunostaining in DIV13 primary neuron cultlures. The percent nuclei that stained positively for each celltype marker are quantified on the right. Three images were analyzed for each marker. **b.** PCA plot of normalized gene expression matrix from ETP and DMSO-treated neurons. Principle component 1 (PC1): 75% variance. Principle component 2 (PC2): 7% variance. **c.** Volcano plot of ETP vs. DMSO contrast. Gray circles: non-significant (ns) transcripts. Red circles: transcripts with FDR adjusted p-value <0.05 and log2 fold change >|0.1|. **d.** Heatmap of differentially expressed genes from ETP vs. DMSO contrast. Columns represent biological replicates. **e.** Quantification of Stage 1 and Stage 2 signature enrichment (from bulk RNAseq) in DMSO and ETP-treated neurons. Each data point represents one biological replicate. **f.** Number of Ccl2 puncta (magenta) are quantified for Rbfox3-positive nuclei (2 or more Rbfox3 puncta within or surrounding the nucleus, yellow), and Rbfox3-negative nuclei. Cyan circles outline nuclei. Each datapoint represents one nucleus. Error bars represent standard error of mean (S.E.M.); ****P<0.0001, ***P<0.001, *P<0.05. Wilcoxon test (e). Student’s t-test (f). Data are representative of two independent experiments (a,f).

**Supplementary Figure 5.**
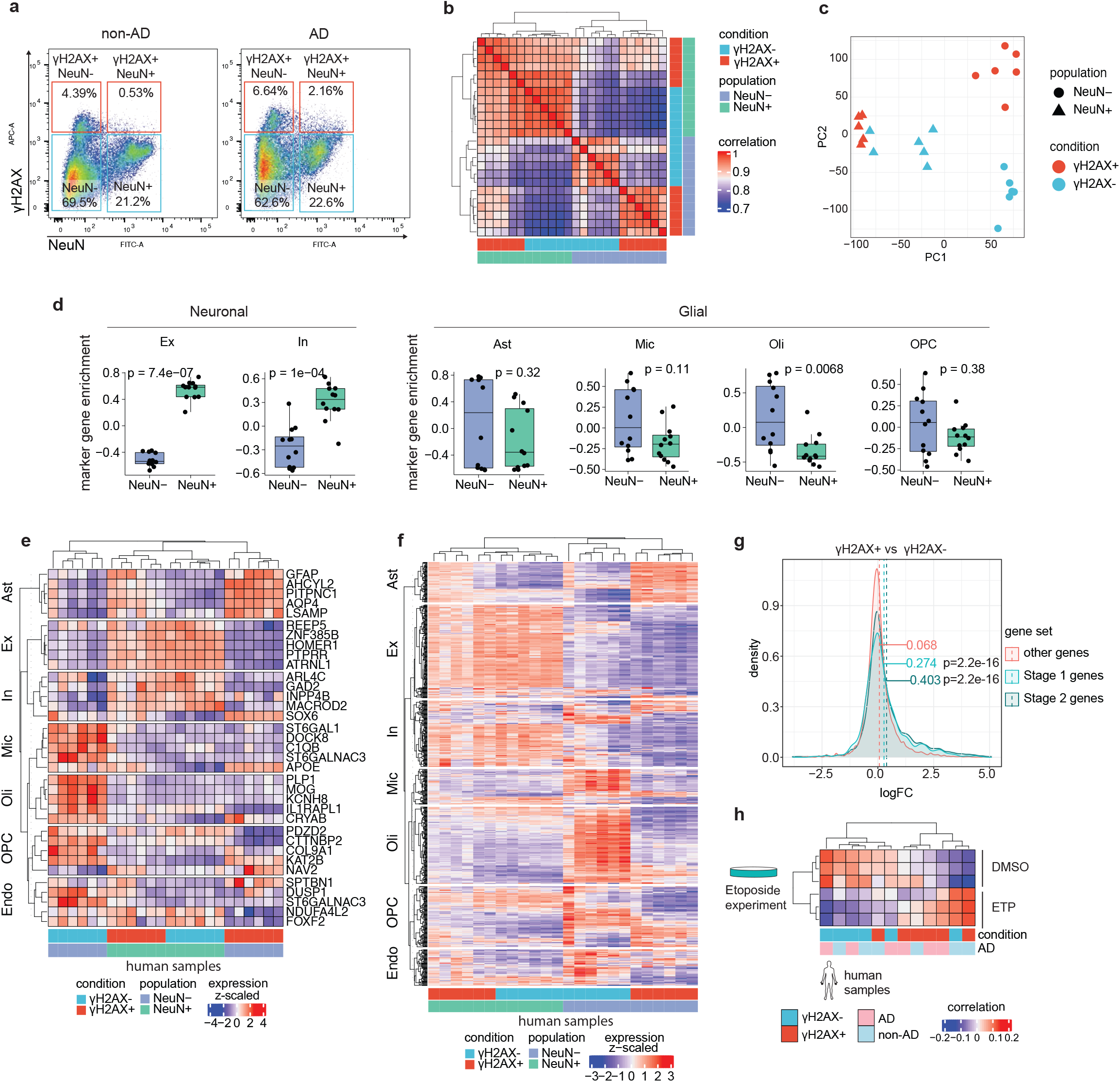
**a.** Flow cytometry dot plot of γH2AX gating for human NeuN+ and NeuN- nuclei. Percent total population is indicated for each gating box. **b.** Correlation heatmap of γH2AX+ and γH2AX-/NeuN+ and NeuN- bulk RNAseq libraries from postmortem temporal cortex. **c.** PCA plot of normalized gene expression from γH2AX+ and γH2AX-/NeuN+ and NeuN- samples. **d.** Quantification of normalized neuronal and glial marker gene expression in NeuN+ and NeuN- samples. Mean gene expression is compared with a Wilcoxon test. Gene set enrichment was calculated using Gene Set Variation Analysis (GSVA). **e.** Gene expression heatmap of top five marker genes for major brain cell types. Each column represents one human sample. **f.** Gene expression heatmap of all marker genes for major brain cell types. Each column represents one human sample. **g.** Distribution of logFC values for Stage 1 genes (light blue), Stage 2 genes (dark blue), and other genes (red). LogFC values are from γH2AX+ vs γH2AX- comparison. Mean LogFC for each gene set are compared with a Wilcoxon test. **h.** Heatmap of ETP and γH2AX+ and γH2AX- human neuronal nuclei transcriptional correlation. Excitatory (Ex), Inihibtory (In), Astrocyte (Ast), Microglia (Mic), Oligodendrocyte (Oli), Oligodendrocyte Precursor Cells (OPC), Endothelial (Endo).

**Supplementary Figure 6.**
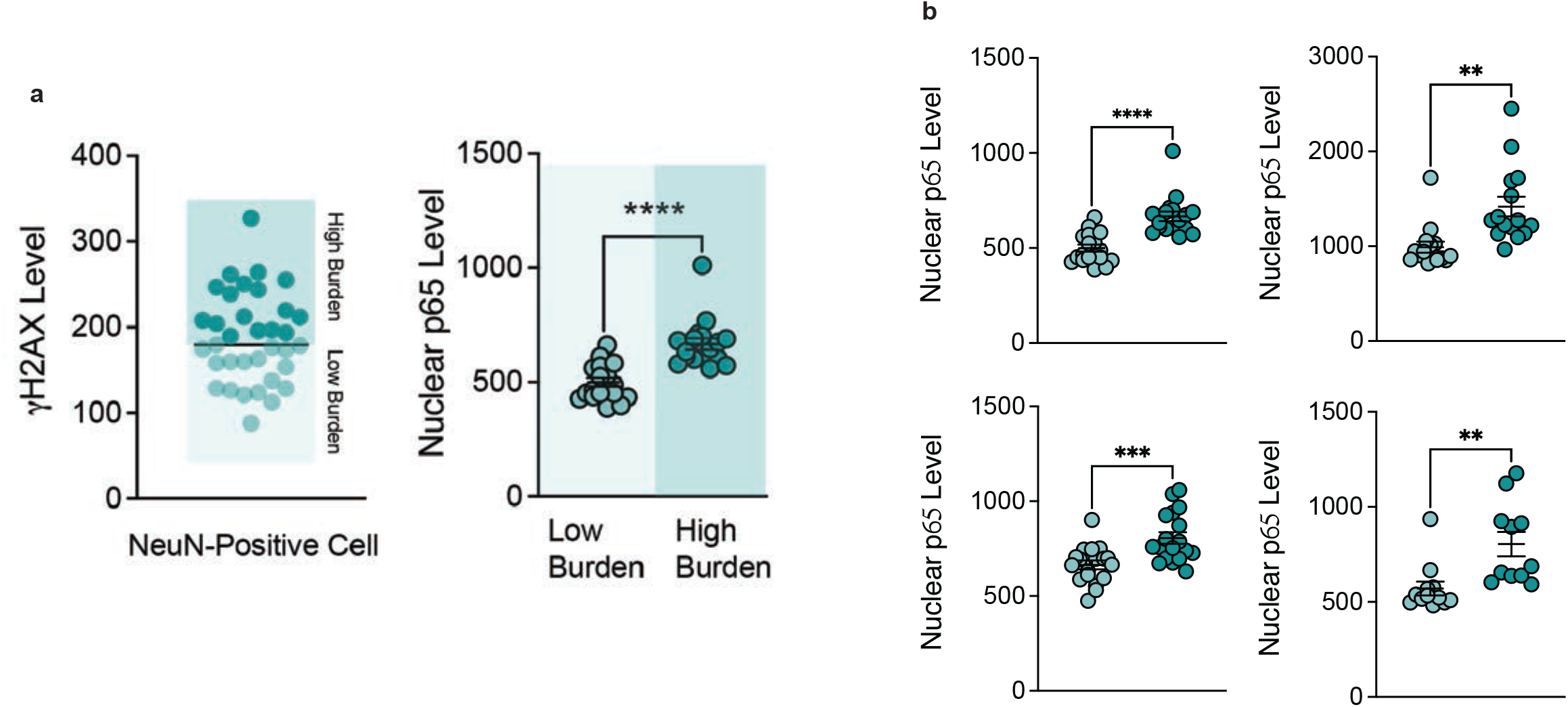
**a.** Schematic for binning γH2AX expression in NeuN-positive cells, and nuclear p65 quanitification. For each individual, the median γH2AX expression in NeuN-positive cells was calculated. Cells were then binned as γH2AX high or γH2AX low based off of median γH2AX expression. Nuclear p65 intensity was then calculated for all cells. **b.** Raw data from the four AD individuals used for analysis. Error bars represent standard error of mean (S.E.M.); ****P<0.0001, ***P<0.001, **P<0.01, *P<0.05, ns not significant. A t-test was performed to compare mean p65 intensity between γH2AX low and γH2AX high cells.

**Supplementary Figure 7.**
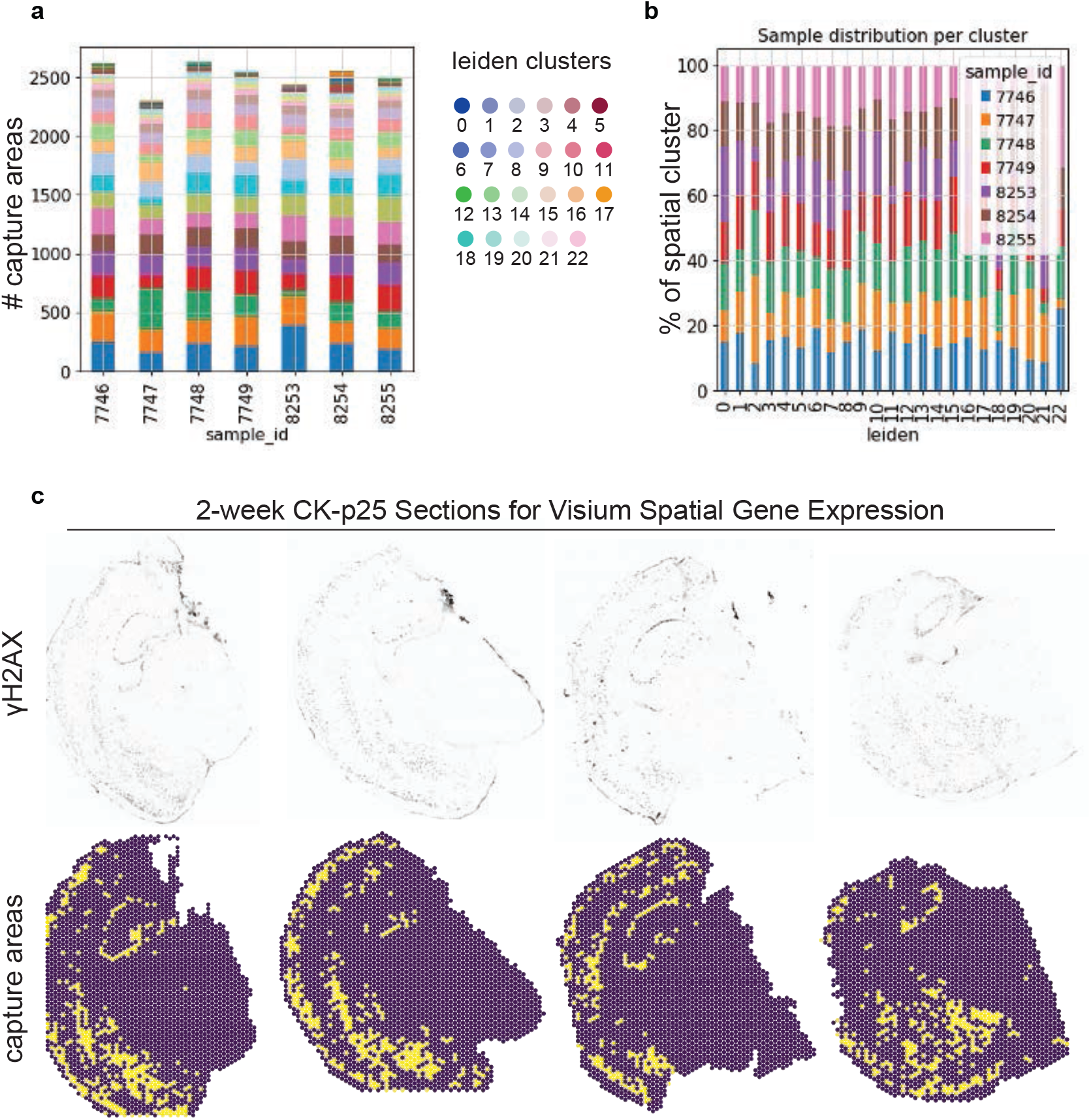
**a.** Distrubution of capture areas amongst spatial clusters for each sample. The y-axis refers to number capture areas. Colors refer to spatial cluster id. **b.** Distribution of samples by spatial cluster. Colors refer to sample id. **c.** Top: γH2AX immunostaining for each CK-p25 section. Bottom: the corresponding capture areas identified as γH2AX-positive are marked in yellow. γH2AX-negative capture areas are marked in purple.

**Supplementary Figure 8.**
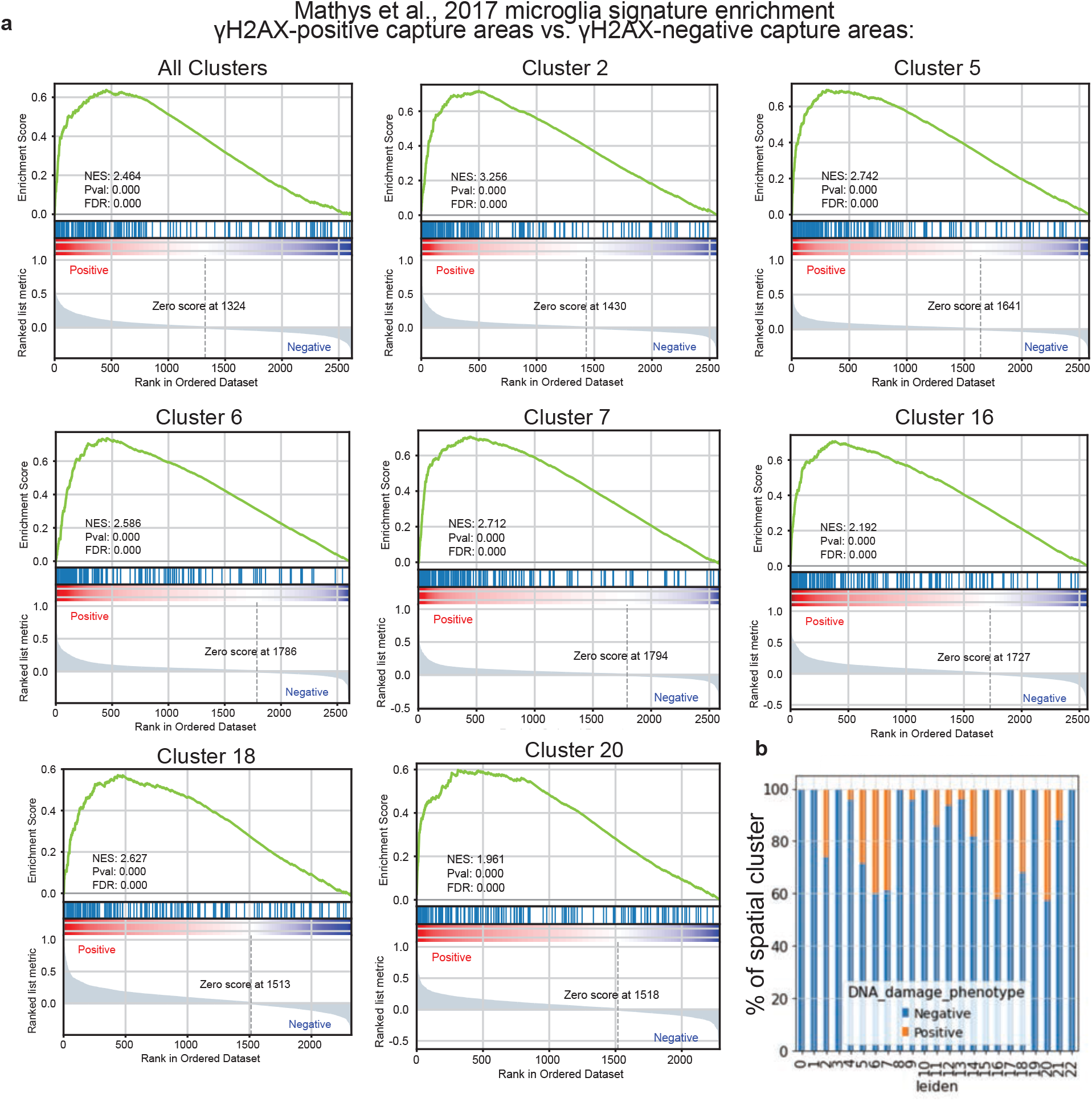
**a.** Gene set enrichment analysis of γH2AX-positive capture area DEGs. DEGs were tested for the enrichment of the reactive microglia signature characterized in Mathys et al., 2019. The analysis was first performed for all γH2AX-posistive capture areas, then for individual clusters comprised of 20% or more γH2AX-positive capture areas. Normalized enrichment score (NES). **b.** Percent γH2AX-positive capture areas by spatial cluster. Orange indicates γH2AX-postive capture areas. Blue indicates γH2AX-negative capture areas.

**Supplementary Figure 9.**
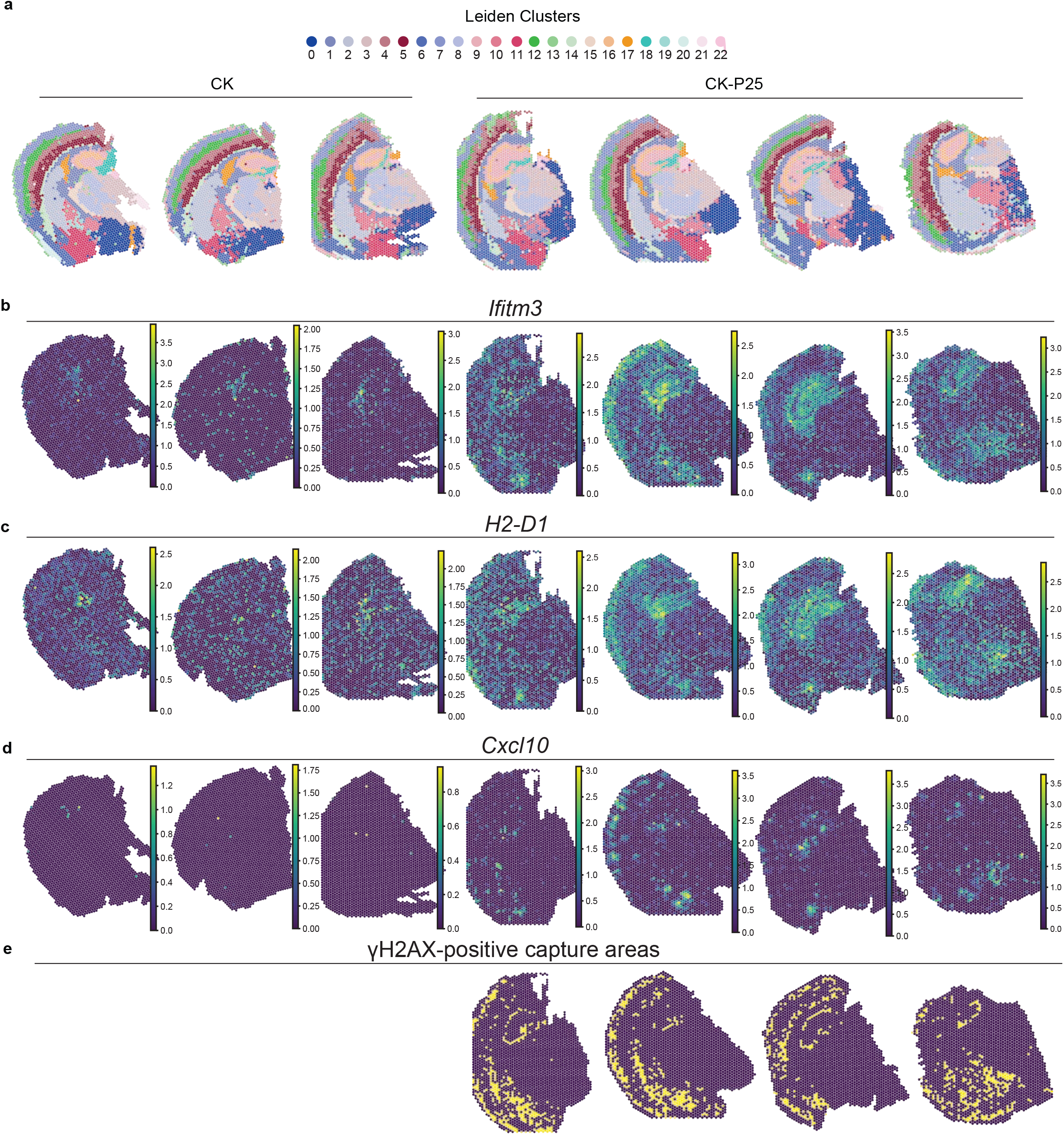
**a.** Distribution of spatial clusters for each sample. **b-d.** Expression of reactive microglia and DSB-bearing neuron genes (b) Ifitm3, (c) H2-D1, and (c) Cxcl10 for each sample. **e.** Distribution of γH2AX-positive capture areas for each CK-p25 sample.

**Supplementary Figure 10.**
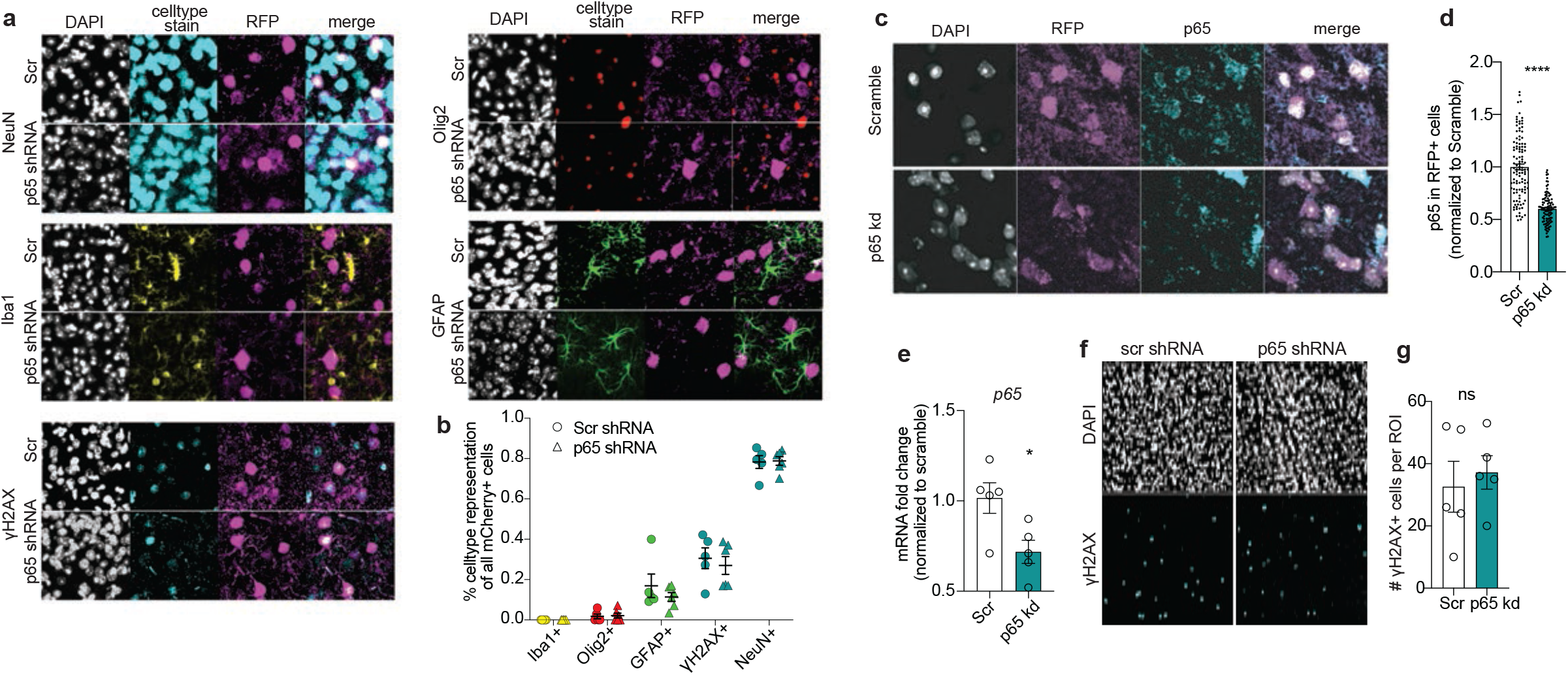
**a.** Representative images of cell type specific immunostaining and RFP immunostaining in scramble and p65kd cortex. Top to bottom, left to right: NeuN=neurons, Iba1=microglia, γH2AX=γH2AX+ cells, Olig2=oligodendrocytes and oligodendrocyte precursor cells, GFAP=astrocytes. **b.** Quantification of cell type distribution of RFP+ cells. Total number of RFP+ cells are quantified per image, then the fraction co-stained for a celltype marker are calculated. Each datapoint represents one mouse. **c.** P65 and RFP immunostaining in scramble and p65kd CK-p25 cortex. **d.** Quantification of p65 mean intensity. Analysis was performed on three animals, two sections each. Scramble n=113 cells, p65kd n=103 cells. **e.** qRT-PCR of p65 in RFP+ NeuN+ nuclei from scramble and shp65-treated CK-p25 mice. **f.** Representative images of γH2AX immunostaining in p65 kd and scramble CK-p25 cortex. **g.** Number of γH2AX+ per image are quantified. Each data point represents one mouse. Error bars represent standard error of mean (S.E.M.); ****P<0.0001, ***P<0.001, **P<0.01, *P<0.05, n.s. not significant. Student’s t-test (d,e,g).

**Supplementary Figure 11.**
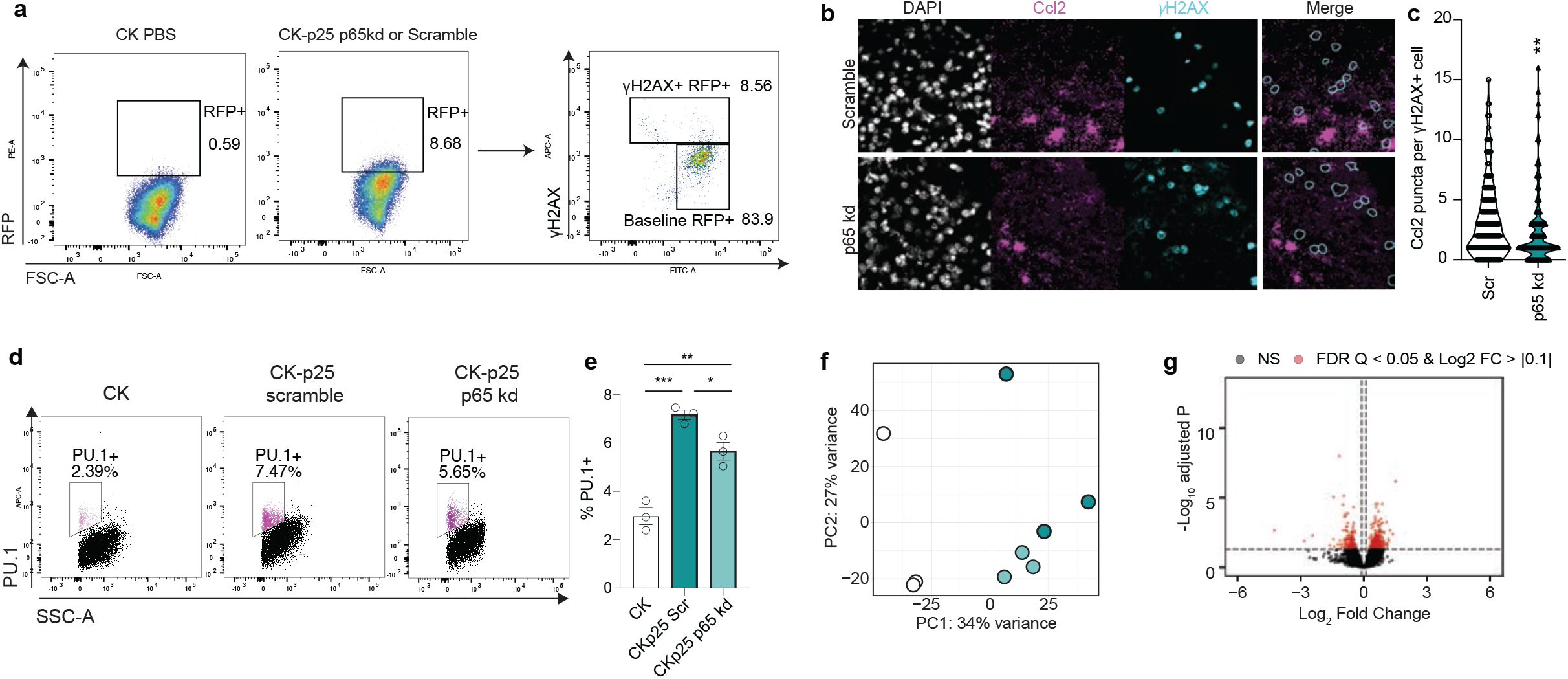
**a.** Sorting schematic for RFP+ γH2AX+ and γH2AX- neurons. 30,000 nuclei were collected for each gate for each animal. **b.** Representative images of Ccl2 RNAscope combined with γH2AX immunofluorescence in Scramble and p65kd cortex. **c.** Quantification of Ccl2 puncta per γH2AX+ cell. Each datapoint represents one cell (n=197 for scramble, n=157 for p65 kd). 20-40 cells were analyzed per mouse. **d.** Sorting schematic for Pu.1+ nuclei for RNA-sequencing. **e.** Quantification of total percent Pu.1+. One datapoint represents one mouse. **f.** PCA plot of normalized gene expression matrix from Pu.1+ bulk RNA-sequencing. **g.** Volcano plot from P65 vs. Scramble contrast. Error bars represent standard error of mean (S.E.M.); ****P<0.0001, ***P<0.001, **P<0.01, *P<0.05, ns not significant. One-way ANOVA followed by Tukey’s test for multiple comparisons (e).

**Supplementary Table 3.**

| ADRC # | Clinical Diagnosis | SEX | PMI | AGE | Braak | Demented? |
| --- | --- | --- | --- | --- | --- | --- |
| 2018 | Control | F | 24 | 92 | II | No |
| 2068 | Control | F | 9 | 79 | II | No |
| 2088 | Control | M | 24 | 65 | I | No |
| 2136 | AD | F | 24 | 87 | VI | Yes |
| 2166 | AD | F | 21 | 71 | VI | Yes |
| 2151 | AD | M | 23 | 81 | VI | Yes |

Clinical Diagnosis = Diagnosis at time of death based on Clinician and Pathology fellow's knowledge SEX F=Female, M=Male
PMI = Post mortem interval (in hours)
AGE = Age at death (in hours)
Braak = Braack stage

**Supplementary Table 4.**
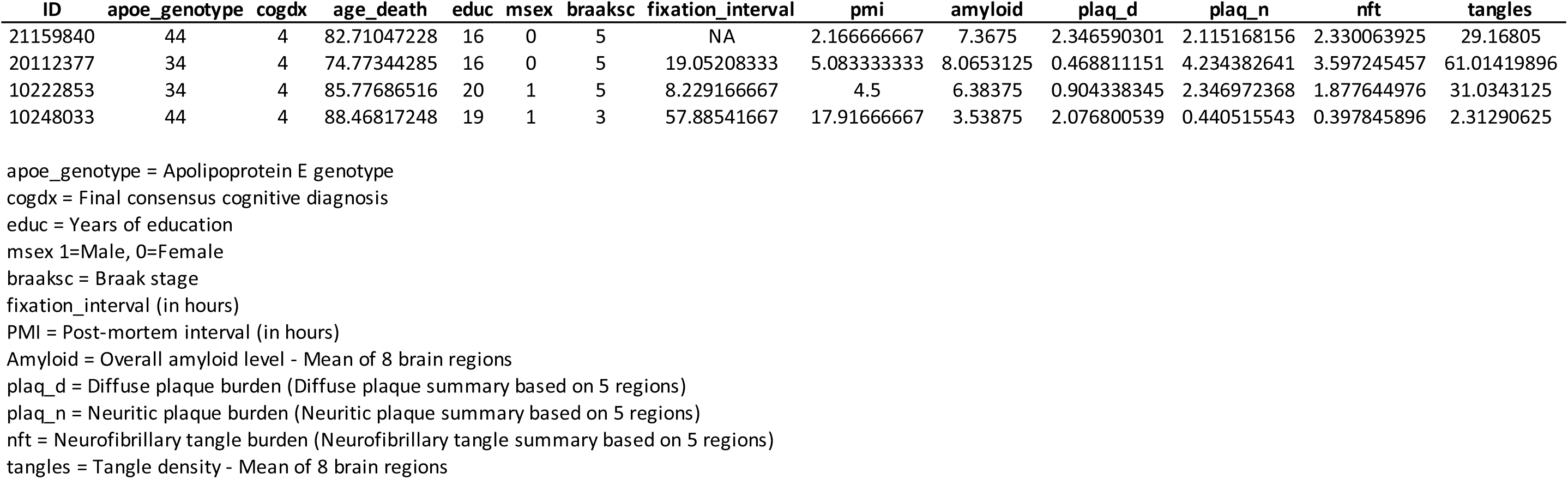

## References

1. Madabhushi, R., Pan, L. & Tsai, L.-H. DNA Damage and Its Links to Neurodegeneration. Neuron 83, 266–282 (2014).

2. Martin, L. J. DNA Damage and Repair: Relevance to Mechanisms of Neurodegeneration. J. Neuropathol. Exp. Neurol. 67, 377–387 (2008).

3. Hoeijmakers, J. H. J. DNA Damage, Aging, and Cancer. N. Engl. J. Med. 361, 1475–1485 (2009).

4. Lu, T. et al. Gene regulation and DNA damage in the ageing human brain. Nature 429, 883–891 (2004).

5. Adamec, E., Vonsattel, J. P. & Nixon, R. A. DNA strand breaks in Alzheimer’s disease. Brain Res. 849, 67–77 (1999).

6. Lovell, M. A. & Markesbery, W. R. Oxidative DNA damage in mild cognitive impairment and late-stage Alzheimer’s disease. Nucleic Acids Res. 35, 7497–7504 (2007).

7. Jacobsen, E., Beach, T., Shen, Y., Li, R. & Chang, Y. Deficiency of the Mre11 DNA repair complex in Alzheimer’s disease brains. Mol. Brain Res. 128, 1–7 (2004).

8. Shanbhag, N. M. et al. Early neuronal accumulation of DNA double strand breaks in Alzheimer’s disease. Acta Neuropathol. Commun. 7, 77 (2019).

9. Kim, D. et al. Deregulation of HDAC1 by p25/Cdk5 in Neurotoxicity. Neuron 60, 803–817 (2008).

10. Sykora, P. et al. DNA polymerase β deficiency leads to neurodegeneration and exacerbates Alzheimer disease phenotypes. Nucleic Acids Res. 43, 943–959 (2015).

11. Dobbin, M. M. et al. SIRT1 collaborates with ATM and HDAC1 to maintain genomic stability in neurons. Nat. Neurosci. 16, 1008–1015 (2013).

12. Suberbielle, E. et al. DNA repair factor BRCA1 depletion occurs in Alzheimer brains and impairs cognitive function in mice. Nat. Commun. 6, 8897 (2015).

13. Suberbielle, E. et al. Physiologic brain activity causes DNA double-strand breaks in neurons, with exacerbation by amyloid-β. Nat. Neurosci. 16, 613–621 (2013).

14. Hayano, M. et al. DNA Break-Induced Epigenetic Drift as a Cause of Mammalian Aging. https://papers.ssrn.com/abstract=3466338 (2019) doi:10.2139/ssrn.3466338.

15. Wu, W. et al. Neuronal enhancers are hotspots for DNA single-strand break repair. Nature 593, 440–444 (2021).

16. Reid, D. A. et al. Incorporation of a nucleoside analog maps genome repair sites in postmitotic human neurons. Science 372, 91–94 (2021).

17. Härtlova, A. et al. DNA Damage Primes the Type I Interferon System via the Cytosolic DNA Sensor STING to Promote Anti-Microbial Innate Immunity. Immunity 42, 332–343 (2015).

18. Dou, Z. et al. Cytoplasmic chromatin triggers inflammation in senescence and cancer. Nature 550, 402–406 (2017).

19. Janssens, S. & Tschopp, J. Signals from within: the DNA-damage-induced NF-κB response. Cell Death Differ. 13, 773–784 (2006).

20. McCool, K. W. & Miyamoto, S. DNA damage-dependent NF-κB activation: NEMO turns nuclear signaling inside out. Immunol. Rev. 246, 311–326 (2012).

21. Rodier, F. et al. Persistent DNA damage signalling triggers senescence-associated inflammatory cytokine secretion. Nat. Cell Biol. 11, 973–979 (2009).

22. Chien, Y. et al. Control of the senescence-associated secretory phenotype by NF-B promotes senescence and enhances chemosensitivity. Genes Dev. 25, 2125–2136 (2011).

23. Klein, R. S. et al. Neuronal CXCL10 Directs CD8+ T-Cell Recruitment and Control of West Nile Virus Encephalitis. J. Virol. 79, 11457–11466 (2005).

24. Préhaud, C., Mégret, F., Lafage, M. & Lafon, M. Virus Infection Switches TLR-3-Positive Human Neurons To Become Strong Producers of Beta Interferon. J. Virol. 79, 12893–12904 (2005).

25. Cruz, J. C., Tseng, H.-C., Goldman, J. A., Shih, H. & Tsai, L.-H. Aberrant Cdk5 Activation by p25 Triggers Pathological Events Leading to Neurodegeneration and Neurofibrillary Tangles. Neuron 40, 471–483 (2003).

26. Mathys, H. et al. Temporal Tracking of Microglia Activation in Neurodegeneration at Single-Cell Resolution. Cell Rep. 21, 366–380 (2017).

27. Cruz, J. C. et al. p25/Cyclin-Dependent Kinase 5 Induces Production and Intraneuronal Accumulation of Amyloid beta In Vivo. J. Neurosci. 26, 10536–10541 (2006).

28. Fischer, A., Sananbenesi, F., Pang, P. T., Lu, B. & Tsai, L.-H. Opposing Roles of Transient and Prolonged Expression of p25 in Synaptic Plasticity and Hippocampus-Dependent Memory. Neuron 48, 825–838 (2005).

29. Gjoneska, E. et al. Conserved epigenomic signals in mice and humans reveal immune basis of Alzheimer’s disease. Nature 518, 365–369 (2015).

30. Mah, L.-J., El-Osta, A. & Karagiannis, T. C. γH2AX: a sensitive molecular marker of DNA damage and repair. Leukemia 24, 679–686 (2010).

31. Caldwell, A. B. et al. Dedifferentiation and neuronal repression define familial Alzheimer’s disease. Sci. Adv. 6, eaba5933 (2020).

32. Mertens, J. et al. Age-dependent instability of mature neuronal fate in induced neurons from Alzheimer’s patients. Cell Stem Cell (2021) doi:10.1016/j.stem.2021.04.004.

33. Yousef, A. et al. Neuron loss and degeneration in the progression of TDP-43 in frontotemporal lobar degeneration. Acta Neuropathol. Commun. 5, 68 (2017).

34. Ünal-Çevik, I., Kılınç, M., Gürsoy-Özdemir, Y., Gurer, G. & Dalkara, T. Loss of NeuN immunoreactivity after cerebral ischemia does not indicate neuronal cell loss: a cautionary note. Brain Res. 1015, 169–174 (2004).

35. McPhail, L. T., McBride, C. B., McGraw, J., Steeves, J. D. & Tetzlaff, W. Axotomy abolishes NeuN expression in facial but not rubrospinal neurons. Exp. Neurol. 185, 182–190 (2004).

36. Wu, K.-L. et al. Loss of Neuronal Protein Expression in Mouse Hippocampus After Irradiation. J. Neuropathol. Exp. Neurol. 69, 272–280 (2010).

37. Collombet, J.-M. et al. Early reduction of NeuN antigenicity induced by soman poisoning in mice can be used to predict delayed neuronal degeneration in the hippocampus. Neurosci. Lett. 398, 337–342 (2006).

38. Subramanian, A. et al. Gene set enrichment analysis: A knowledge-based approach for interpreting genome-wide expression profiles. Proc. Natl. Acad. Sci. 102, 15545–15550 (2005).

39. Yang, Y., Geldmacher, D. S. & Herrup, K. DNA Replication Precedes Neuronal Cell Death in Alzheimer’s Disease. J. Neurosci. 21, 2661–2668 (2001).

40. Busser, J., Geldmacher, D. S. & Herrup, K. Ectopic Cell Cycle Proteins Predict the Sites of Neuronal Cell Death in Alzheimer’s Disease Brain. J. Neurosci. 18, 2801–2807 (1998).

41. Sokolova, A. et al. Monocyte Chemoattractant Protein-1 Plays a Dominant Role in the Chronic Inflammation Observed in Alzheimer’s Disease. Brain Pathol. 19, 392–398 (2009).

42. El Khoury, J., et al. Ccr2 deficiency impairs microglial accumulation and accelerates progression of Alzheimer-like disease. Nat. Med. 13, 432–438 (2007).

43. Chai, Q., She, R., Huang, Y. & Fu, Z. F. Expression of Neuronal CXCL10 Induced by Rabies Virus Infection Initiates Infiltration of Inflammatory Cells, Production of Chemokines and Cytokines, and Enhancement of Blood-Brain Barrier Permeability. J. Virol. 89, 870–876 (2014).

44. Conductier, G., Blondeau, N., Guyon, A., Nahon, J.-L. & Rovère, C. The role of monocyte chemoattractant protein MCP1/CCL2 in neuroinflammatory diseases. J. Neuroimmunol. 224, 93–100 (2010).

45. Chen, E. Y. et al. Enrichr: interactive and collaborative HTML5 gene list enrichment analysis tool. BMC Bioinformatics 14, 128 (2013).

46. Kuleshov, M. V. et al. Enrichr: a comprehensive gene set enrichment analysis web server 2016 update. Nucleic Acids Res. 44, W90–97 (2016).

47. Xie, Z. et al. Gene Set Knowledge Discovery with Enrichr. Curr. Protoc. 1, e90 (2021).

48. Trapnell, C. et al. The dynamics and regulators of cell fate decisions are revealed by pseudotemporal ordering of single cells. Nat. Biotechnol. 32, 381–386 (2014).

49. Qiu, X. et al. Reversed graph embedding resolves complex single-cell trajectories. Nat. Methods 14, 979–982 (2017).

50. Cao, J. et al. The single-cell transcriptional landscape of mammalian organogenesis. Nature 566, 496–502 (2019).

51. Wang, C. et al. Gain of toxic apolipoprotein E4 effects in human iPSC-derived neurons is ameliorated by a small-molecule structure corrector. Nat. Med. 24, 647–657 (2018).

52. Dunphy, G. et al. Non-canonical Activation of the DNA Sensing Adaptor STING by ATM and IFI16 Mediates NF-κB Signaling after Nuclear DNA Damage. Mol. Cell 71, 745–760.e5 (2018).

53. Mathys, H. et al. Single-cell transcriptomic analysis of Alzheimer’s disease. Nature 570, 332–337 (2019).

54. Variable Details | RADC. https://www.radc.rush.edu/docs/var/detail.htm?category=Pathology&subcategory=Alzheimer%27s+disease&variable=gpath.

55. Chan, K. Y. et al. Engineered AAVs for efficient noninvasive gene delivery to the central and peripheral nervous systems. Nat. Neurosci. 20, 1172–1179 (2017).

56. Chang, A. L. et al. CCL2 Produced by the Glioma Microenvironment Is Essential for the Recruitment of Regulatory T Cells and Myeloid-Derived Suppressor Cells. Cancer Res. 76, 5671–5682 (2016).

57. Semple, B. D., Bye, N., Rancan, M., Ziebell, J. M. & Morganti-Kossmann, M. C. Role of CCL2 (MCP-1) in traumatic brain injury (TBI): evidence from severe TBI patients and CCL2−/− mice. J. Cereb. Blood Flow Metab. Off. J. Int. Soc. Cereb. Blood Flow Metab. 30, 769–782 (2010).

58. Galimberti, D. et al. Serum MCP-1 levels are increased in mild cognitive impairment and mild Alzheimer’s disease. Neurobiol. Aging 27, 1763–1768 (2006).

59. Bradburn, S. et al. Dysregulation of C-X-C motif ligand 10 during aging and association with cognitive performance. Neurobiol. Aging 63, 54–64 (2018).

60. Zhang, B. et al. Integrated Systems Approach Identifies Genetic Nodes and Networks in Late-Onset Alzheimer’s Disease. Cell 153, 707–720 (2013).

61. Keren-Shaul, H. et al. A Unique Microglia Type Associated with Restricting Development of Alzheimer’s Disease. Cell 169, 1276–1290.e17 (2017).

62. Alzheimer Disease Genetics Consortium (ADGC), et al. Genetic meta-analysis of diagnosed Alzheimer’s disease identifies new risk loci and implicates Aβ, tau, immunity and lipid processing. Nat. Genet. 51, 414–430 (2019).

63. Nott, A. et al. Brain cell type–specific enhancer–promoter interactome maps and disease **-** risk association. Science 366, 1134–1139 (2019).

64. Ramamurthy, E. et al. Cell type-specific histone acetylation profiling of Alzheimer’s Disease subjects and integration with genetics. 2020.03.26.010330 https://www.biorxiv.org/content/10.1101/2020.03.26.010330v1 (2020) doi:10.1101/2020.03.26.010330.

65. Shi, Y. et al. Microglia drive APOE-dependent neurodegeneration in a tauopathy mouse model. J. Exp. Med. 216, 2546–2561 (2019).

66. Spangenberg, E. et al. Sustained microglial depletion with CSF1R inhibitor impairs parenchymal plaque development in an Alzheimer’s disease model. Nat. Commun. 10, 3758 (2019).

67. Spangenberg, E. E. et al. Eliminating microglia in Alzheimer’s mice prevents neuronal loss without modulating amyloid-β pathology. Brain 139, 1265–1281 (2016).

68. Song, W. M. & Colonna, M. The identity and function of microglia in neurodegeneration. Nat. Immunol. 19, 1048–1058 (2018).

69. Simon, M. et al. LINE1 Derepression in Aged Wild-Type and SIRT6-Deficient Mice Drives Inflammation. Cell Metab. 29, 871–885.e5 (2019).

70. Lee, H. et al. Cell Type-Specific Transcriptomics Reveals that Mutant Huntingtin Leads to Mitochondrial RNA Release and Neuronal Innate Immune Activation. Neuron 107, 891–908.e8 (2020).

71. Wyss-Coray, T. & Rogers, J. Inflammation in Alzheimer Disease—A Brief Review of the Basic Science and Clinical Literature. Cold Spring Harb. Perspect. Med. 2, (2012).

72. Kaltschmidt, B. et al. NF-kappaB regulates spatial memory formation and synaptic plasticity through protein kinase A/CREB signaling. Mol. Cell. Biol. 26, 2936–2946 (2006).

73. Fridmacher, V. et al. Forebrain-Specific Neuronal Inhibition of Nuclear Factor-κB Activity Leads to Loss of Neuroprotection. J. Neurosci. 23, 9403–9408 (2003).

74. Mahad, D. et al. Modulating CCR2 and CCL2 at the blood–brain barrier: relevance for multiple sclerosis pathogenesis. Brain 129, 212–223 (2006).

75. Dimitrijevic, O. B., Stamatovic, S. M., Keep, R. F. & Andjelkovic, A. V. Effects of the Chemokine CCL2 on Blood–Brain Barrier Permeability during Ischemia–Reperfusion Injury. J. Cereb. Blood Flow Metab. 26, 797–810 (2006).

76. Naert, G. & Rivest, S. CC Chemokine Receptor 2 Deficiency Aggravates Cognitive Impairments and Amyloid Pathology in a Transgenic Mouse Model of Alzheimer’s Disease. J. Neurosci. 31, 6208–6220 (2011).

77. Yamamoto, M. et al. Overexpression of Monocyte Chemotactic Protein-1/CCL2 in β-Amyloid Precursor Protein Transgenic Mice Show Accelerated Diffuse β-Amyloid Deposition. Am. J. Pathol. 166, 1475–1485 (2005).

78. Krauthausen, M. et al. CXCR3 promotes plaque formation and behavioral deficits in an Alzheimer’s disease model. J. Clin. Invest. 125, 365–378 (2015).

79. Young, K. & Morrison, H. Quantifying Microglia Morphology from Photomicrographs of Immunohistochemistry Prepared Tissue Using ImageJ. J. Vis. Exp. 57648 (2018) doi:10.3791/57648.

## Methods References

1. Kolberg, Liis, Uku Raudvere, Ivan Kuzmin, Jaak Vilo, and Hedi Peterson. “Gprofiler2 -- an R Package for Gene List Functional Enrichment Analysis and Namespace Conversion Toolset g:Profiler.” F1000Research 9 (November 17, 2020): 709. https://doi.org/10.12688/f1000research.24956.2.

2. Liao, Yang, Gordon K. Smyth, and Wei Shi. “The R Package Rsubread Is Easier, Faster, Cheaper and Better for Alignment and Quantification of RNA Sequencing Reads.” Nucleic Acids Research 47, no. 8 (May 7, 2019): e47. https://doi.org/10.1093/nar/gkz114.

3. Mohammadi, Shahin, Jose Davila-Velderrain, and Manolis Kellis. “A Multiresolution Framework to Characterize Single-Cell State Landscapes.” Nature Communications 11, no. 1 (October 26, 2020): 5399. https://doi.org/10.1038/s41467-020-18416-6.

4. Robinson, Mark D., Davis J. McCarthy, and Gordon K. Smyth. “EdgeR: A Bioconductor Package for Differential Expression Analysis of Digital Gene Expression Data.” Bioinformatics (Oxford, England) 26, no. 1 (January 1, 2010): 139–40. https://doi.org/10.1093/bioinformatics/btp616.

5. Bennett, David A., Aron S. Buchman, Patricia A. Boyle, Lisa L. Barnes, Robert S. Wilson, and Julie A. Schneider. “Religious Orders Study and Rush Memory and Aging Project.” Journal of Alzheimer’s Disease: JAD 64, no. s1 (2018): S161–89. https://doi.org/10.3233/JAD-179939.

6. Swiech, Lukasz, Matthias Heidenreich, Abhishek Banerjee, Naomi Habib, Yinqing Li, John Trombetta, Mriganka Sur, and Feng Zhang. “In Vivo Interrogation of Gene Function in the Mammalian Brain Using CRISPR-Cas9.” Nature Biotechnology 33, no. 1 (January 2015): 102–6. https://doi.org/10.1038/nbt.3055.

7. Dobin, Alexander, Carrie A. Davis, Felix Schlesinger, Jorg Drenkow, Chris Zaleski, Sonali Jha, Philippe Batut, Mark Chaisson, and Thomas R. Gingeras. “STAR: Ultrafast Universal RNA-Seq Aligner.” Bioinformatics (Oxford, England) 29, no. 1 (January 1, 2013): 15–21. https://doi.org/10.1093/bioinformatics/bts635.

8. Anders, Simon, Paul Theodor Pyl, and Wolfgang Huber. “HTSeq--a Python Framework to Work with High-Throughput Sequencing Data.” Bioinformatics (Oxford, England) 31, no. 2 (January 15, 2015): 166–69. https://doi.org/10.1093/bioinformatics/btu638.

9. Wolf, F. Alexander, Philipp Angerer, and Fabian J. Theis. “SCANPY: Large-Scale Single-Cell Gene Expression Data Analysis.” Genome Biology 19, no. 1 (February 6, 2018): 15. https://doi.org/10.1186/s13059-017-1382-0.

10. De Jager, Philip L., Yiyi Ma, Cristin McCabe, Jishu Xu, Badri N. Vardarajan, Daniel Felsky, Hans-Ulrich Klein, et al. “A Multi-Omic Atlas of the Human Frontal Cortex for Aging and Alzheimer’s Disease Research.” Scientific Data 5, no. 1 (August 7, 2018): 180142. https://doi.org/10.1038/sdata.2018.142.

11. Storey, John D., Andrew J. Bass, Alan Dabney, David Robinson, and Gregory Warnes. Qvalue: Q-Value Estimation for False Discovery Rate Control (version 2.22.0). Bioconductor version: Release (3.12), 2021. https://doi.org/10.18129/B9.bioc.qvalue.

12. Raudvere, Uku, Liis Kolberg, Ivan Kuzmin, Tambet Arak, Priit Adler, Hedi Peterson, and Jaak Vilo. “G:Profiler: A Web Server for Functional Enrichment Analysis and Conversions of Gene Lists (2019 Update).” Nucleic Acids Research 47, no. W1 (July 2, 2019): W191–98. https://doi.org/10.1093/nar/gkz369.

13. Jassal, Bijay, Lisa Matthews, Guilherme Viteri, Chuqiao Gong, Pascual Lorente, Antonio Fabregat, Konstantinos Sidiropoulos, et al. “The Reactome Pathway Knowledgebase.” Nucleic Acids Research 48, no. D1 (January 8, 2020): D498–503. https://doi.org/10.1093/nar/gkz1031.

14. Ligtenberg, W. “Reactome.Db: A Set of Annotation Maps for Reactome Assembled Using Data from Reactome.” Bioconductor, 2019. http://bioconductor.org/packages/reactome.db/.

15. Mathys, Hansruedi, Jose Davila-Velderrain, Zhuyu Peng, Fan Gao, Shahin Mohammadi, Jennie Z. Young, Madhvi Menon, et al. “Single-Cell Transcriptomic Analysis of Alzheimer’s Disease.” Nature 570, no. 7761 (June 2019): 332–37. https://doi.org/10.1038/s41586-019-1195-2.

